# Oxytocin mediates the acquisition and strategy formation of cooperation in rats

**DOI:** 10.64898/2026.03.02.708935

**Authors:** Yongqin Lin, Lei Wei, Qingxiu Wang, Zuoren Wang

## Abstract

Cooperative behaviors are widespread in nature and play fundamental roles throughout animal lifespans. However, the learning processes and strategy formation underlying cooperation remain largely unknown. Oxytocin (OXT) is a neuropeptide that regulates both social and non-social behaviors, yet its roles in modulating cooperative behaviors remain to be fully elucidated. To investigate these questions, we developed a temporal coordination-based cooperative paradigm for rats and manipulated the OXT system during training. Through progressive cooperative training, rats acquired cooperation and efficiently performed the task via social interaction. This learning process was accompanied by increased OXT release in social learning-associated brain regions, whereas OXT knockout impaired learning speed. Furthermore, rats displayed different cooperative strategies, with the communication-based strategy gradually increasing and becoming predominant during the cooperative training. Both OXT knockout and inhibition of OXT-ergic neurons in the paraventricular nucleus (PVN) of the hypothalamus during learning reduced the adoption of communication-based strategy. Together, our results demonstrate that OXT is necessary for both the acquisition and the strategy formation of cooperation in rats, thereby facilitating mutual rewards. These findings provide new insights into the neural mechanisms by which OXT regulates social behaviors and offer potential therapeutic implications for neuropsychiatric disorders involving social deficits, such as autism spectrum disorder and social anxiety disorder.

## INTRODUCTION

In nature, cooperation is a powerful strategy that enhances individual survival and reproduction. Cooperative hunting and defense improve foraging efficiency and survival ^1,2^, while cooperative courtship and breeding enhance mating opportunities and offspring success ^3,4^. As cooperative behaviors provide individuals with positive payoffs or direct fitness benefits, they continue to evolve ^5^. According to behavioral ecologists, cooperation is defined as an outcome in which two or more interacting individuals engage in joint actions, incurring costs to obtain mutual rewards ^6^. To uncover its physiological mechanisms, diverse paradigms have been developed across species. In humans, these paradigms primarily derive from game theory-based economic exchange games ^7,8^. For nonhuman primates, cooperative paradigms include game-theory-based tasks ^9,10^ and non-game-theory-based tasks that require simultaneous actions, such as pulling handles ^11,12^, pulling ropes ^13,14^, or pushing buttons ^15^. Similar rope-pulling tasks have also been applied to species ranging from Asian elephants (*Elephas maximus*) and bottlenose dolphins (*Tursiops truncatus*) to wolves (*Canis Lupus*) ^16–18^.

In rodents, cooperative paradigms were first established in rats (*Rattus norvegicus*) ^19,20^ and later extended to mice (*Mus musculus*) ^21–25^. Tasks such as coordinated shuttling ^26,27^, simultaneous nose-poking ^25,28^, lever-pressing ^29^, and climbing ^30^ have demonstrated that rats can acquire cooperation through cooperative training. Notably, in the simultaneous nose-poking task, rats achieve a success rate of 50–60% ^25,31^, significantly higher than the approximately 20–30% observed in mice ^25,32^. However, despite these advances, the learning processes underlying rodent cooperation remain poorly characterized, particularly regarding the dynamics of temporal coordination, social interactions, and cooperative strategies. Since many cooperative behaviors are learned rather than instinctive ^33^, dissecting this learning process is essential for identifying the behavioral factors that promote cooperation. Moreover, levels of cooperation vary across species ^1,5,34^, spanning categories such as similarity, synchrony, coordination, and collaboration ^1^. Therefore, classifying cooperative strategies and elucidating their formation process could illuminate both species-specific cooperation levels and the evolutionary pathways of cooperative behavior.

Oxytocin (OXT) is a neuropeptide primarily synthesized and released by oxytocinergic neurons in the paraventricular nucleus (PVN) and supraoptic nucleus (SON) of the hypothalamus ^35,36^. First identified for its roles in parturition and lactation ^37,38^, OXT has been revealed by subsequent studies to be involved in regulating diverse social and non-social behaviors ^35,39,40^. Regarding social behaviors, OXT modulates social recognition (or learning) ^41,42^, social preference ^43,44^, and prosocial behaviors such as empathy ^45–47^, consolation ^48,49^, and rescue-like behaviors ^50,51^. Beyond these relatively innate behaviors, human studies indicate that OXT also modulates cooperative behaviors, with a moderate positive effect size ^52,53^. For instance, intranasal OXT administration has been shown to promote in-group trust and cooperation ^54,55^. In animals, cooperative behaviors are accompanied by increased urinary OXT levels ^56–58^, and exogenous OXT has been shown to enhance cooperative responses in species such as meerkats (*Suricata suricatta*), common vampire bats (*Desmodus rotundus*), and rats ^59–61^. Nevertheless, the temporal dynamics of OXT release during cooperation, as well as its specific modulatory mechanisms, remain unclear.

Here, to investigate how OXT regulates the learning process and strategy formation of cooperative behavior, we developed a temporal coordination-based cooperative paradigm for rats and manipulated the OXT system during training. We showed that wild-type rats acquired cooperation rapidly, whereas OXT knockout rats exhibited impaired learning speed. During cooperative training, rats displayed different cooperative strategies, with the communication-based strategy gradually increasing and eventually becoming predominant. Both OXT knockout and inhibition of PVN OXT-ergic neurons reduced the adoption of communication-based strategy. These findings highlight the critical role of OXT in social learning and the formation of cooperative strategies.

## RESULTS

### Acquisition process of cooperation in paired rats

To investigate the learning process of cooperation in rats, paired rats (dyads) were trained in a temporal coordination-based cooperative task ^25^ that required simultaneous nose-poking to receive a water reward (Figure 1A). Dyads first underwent pre-cooperation training to learn to nose-poke individually for water rewards (Stage 1; Figure 1B). Subsequently, they underwent progressive cooperative training to learn to cooperate with their partners (Stage 2; Figure 1B). In this cooperative task, a water reward was delivered to each rat only if both rats nose-poked within a defined time window (success trial; Supplementary Figure S1A, top). Failure to meet this criterion resulted in no reward (failure trial; Supplementary Figure S1A, bottom). According to the lengths of the cooperative time windows (3 s, 2 s, and 1 s), the cooperative task was divided into three consecutive training phases: 3 days of 3-second cooperation (3sCo), 3 days of 2-second cooperation (2sCo), and 12 days of 1-second cooperation (1sCo; Figure 1B).

**Figure 1.**
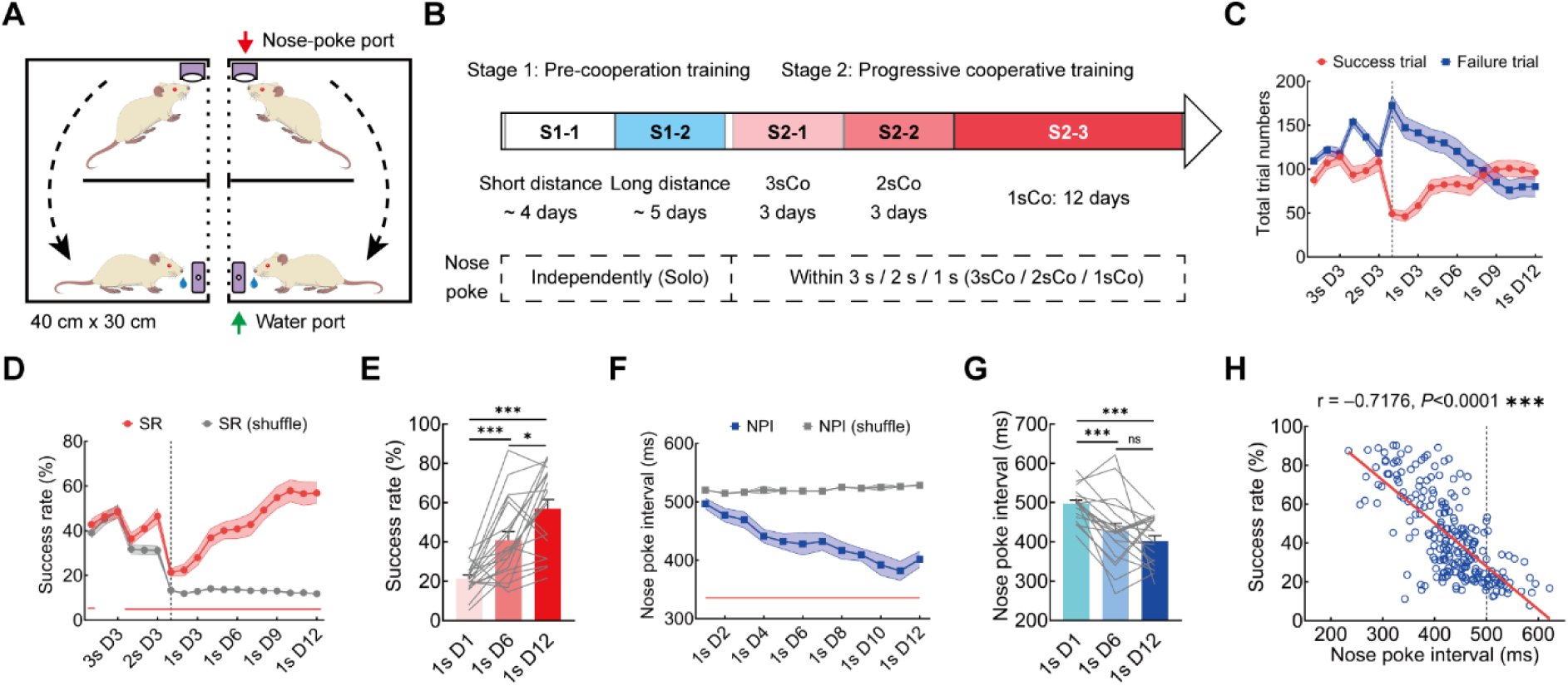
Behavioral performance of paired rats (dyads) during cooperative training. A: Schematic of the behavioral apparatus and task process. Each cage contains a nose-poke port (red arrow), a water port (green arrow) and a 20-cm divider, with middle dashed lines indicating the rectangular perforations. During the task, rats nose-poke (independently or simultaneously) to receive a water reward. B: Schematic of the training process for the cooperative task, which includes pre-cooperation training (Stage 1) and progressive cooperative training (Stage 2); 3sCo, 2sCo, and 1sCo are abbreviations for 3-, 2-, and 1-second cooperation, respectively. C: Total trial numbers of success trials and failure trials for dyads during cooperative training (*n*=20). D: Success rate (SR) of dyads during cooperative training (*n*=20). Gray line indicates the shuffled control of SR. E: Comparisons of SR among the D1, D6, and D12 of 1sCo (D1 vs. D6, *P*=0.0006; D1 vs. D12, *P*<0.0001; D6 vs. D12, *P*=0.0114). F: Nose-poke interval (NPI) of dyads during 1sCo training (*n*=20). Gray line indicates the shuffled control of NPI. G: Comparisons of NPI among the D1, D6, and D12 of 1sCo (D1 vs. D6, *P*=0.0008; D1 vs. D12, *P*<0.0001; D6 vs. D12, *P*=0.4809). H: Linear regression of NPI versus SR during 1sCo (all sessions; *n*=234, *P*<0.0001). “r” is the Pearson correlation coefficient; the vertical dashed line indicates 500 ms NPI. In C and D, vertical dashed lines indicate D1 of 1sCo. In D and F, red lines at the bottom indicate that SR/NPI is statistically different from its shuffled control. Data are presented as mean ± SEM. Statistics: two-tailed paired *t*-test (D and F), repeated measures one-way ANOVA (E and G), and F-test (H). ns: Not significant; *: *P*<0.05; **: *P*<0.01; ***: *P*<0.001.

Throughout cooperative training, dyads exhibited an increase in success trials and a progressive decrease in failure trials during the 1sCo phase (Figure 1C). Moreover, the ratios of each rat initiating nose-poking were approximately equal for both trial types (Supplementary Figure S1B). We further calculated the success rate (SR, success trials/total trials), which gradually increased to nearly 60% during the 1sCo phase (21.42% to 56.87%; Figure 1D, E; red line), significantly exceeding that of the shuffled control (Figure 1D; gray line). These results suggest that dyads were able to acquire cooperation through progressive cooperative training. Behavioral events were also quantified (Supplementary Figure S1C, D). During the 1sCo phase, the nose-poke latency gradually increased, while water lick latency (WLL) and water lick duration (WLD) remained stable (Supplementary Figure S1D).

During both the 3sCo and 2sCo phases, the nose-poke interval (NPI) of dyads remained at their respective chance levels (1 500 ms and 1 000 ms; Supplementary Figure S1E). In contrast, during the 1sCo phase, NPI decreased significantly from the initial chance level of 500 ms to roughly 400 ms (496.83 ms to 401.68 ms; Figure 1F, G). Notably, correlation analysis revealed a strong negative relationship between NPI and SR during the 1sCo phase (r = −0.72; Figure 1H), indicating that better temporal coordination predicted higher cooperation success. Furthermore, 18 days of 3sCo training failed to induce comparable improvements in cooperative performance (Supplementary Figure S1F–H). When well-trained dyads from 3sCo were directly transferred to 1sCo, their SR decreased significantly (Supplementary Figure S1I). Together, these results demonstrate that progressive cooperative training enables rats to acquire cooperation and efficiently perform the task.

### Rats actively await and interact with partners to achieve efficient cooperation

Before initiating nose-poking, rats frequently engaged in social interactions, such as approach behavior ^28^, in which rats might wait for partners or actively interact. To characterize these interactions during cooperative training, we used DeepLabCut ^62,63^, a toolbox for tracking animal behaviors with deep learning, to extract the spatial coordinates of key body parts (nose and ears) from behavioral videos (Supplementary Figure S2A–C). These data were then aligned with behavioral events for joint analyses. Two spatial zones and dual time windows were defined: the nose-poke area (NPA) and water lick area (WLA; Figure 2A), and the waiting time window and social time window (Figure 2B). For success trials, the waiting time window was the interval from one rat entering the NPA until its partner entered the corresponding NPA, and the social time window was the interval from both rats being in the NPA until the first nose-poking occurred (Figure 2B).

**Figure 2.**
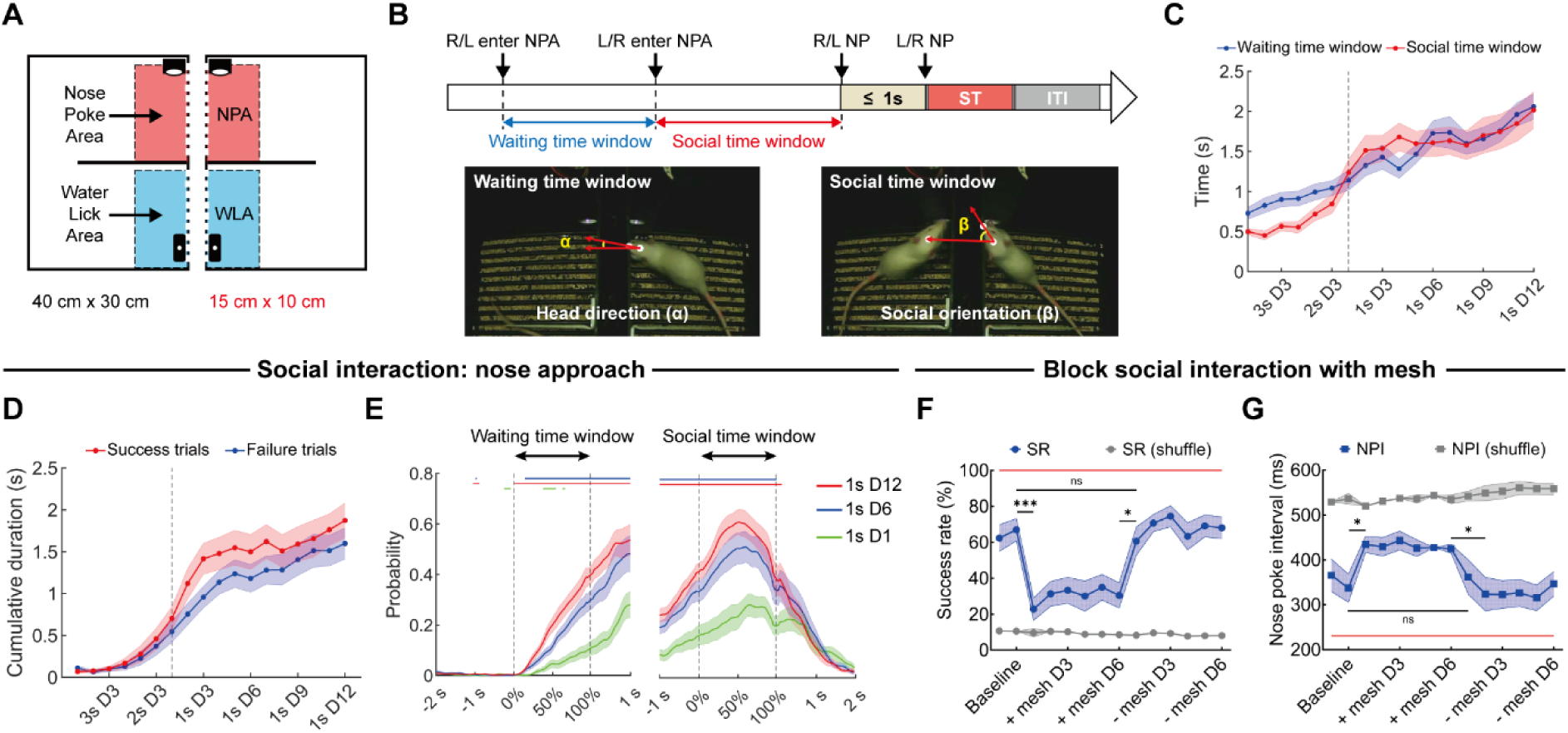
Social interaction between dyads during cooperative training. A: Schematic of the nose-poke area (NPA, red) and the water lick area (WLA, blue). B: Schematic of the waiting time window and social time window during success trials (top); NP, ST, and ITI denote nose-poke, success trial, and inter-trial interval, respectively. Schematics of head direction (α) in the waiting time window (bottom-left) and social orientation (β) in the social time window (bottom-right). C: Changes in the waiting time window (blue) and social time window (red) of dyads during cooperative training (*n*=20). D: Changes in the cumulative duration of nose approach for success trials (red) and failure trials (blue) during cooperative training (*n*=20). E: Probability of nose approach during waiting time window (left) and social time window (right, *n*=20). F: Success rate (SR) of dyads after addition and removal of a metal mesh (blue line, *n*=9; Baseline D2 vs. +mesh D1, *P*=0.0005; +mesh D6 vs. – mesh D1, *P*=0.0129; Baseline D2 vs. -mesh D1, *P*=0.3336). G: Nose-poke interval (NPI) of dyads after addition and removal of a metal mesh (blue line, *n*=9; Baseline D2 vs. +mesh D1, *P*=0.0367; +mesh D6 vs. –mesh D2, *P*=0.0364; Baseline D2 vs. -mesh D1, *P*=0.1217). In C and D, vertical dashed lines indicate D1 of 1sCo. In E, lines at the top indicate statistical significance (*P*<0.05): 1s D1 vs. 1s D6 (blue), 1s D1 vs. 1s D12 (red), and 1s D6 vs. 1s D12 (green). In F and G, gray lines indicate the shuffling control for SR/NPI; red lines at the top/bottom indicate that SR/NPI is statistically different from its shuffled control. Data are presented as mean ± SEM. Statistics: one-way ANOVA (E) and two-tailed paired *t*-test (F–G). ns: Not significant; *: *P*<0.05; **: *P*<0.01; ***: *P*<0.001.

Joint analyses revealed that both the waiting time window (0.73 s to 2.06 s) and the social time window (0.50 s to 2.02 s) increased progressively during cooperative training (Figure 2C; Supplementary Figure S2D), mirroring the increase in the nose-poke latency (Supplementary Figure S1D). Furthermore, the proportion of trials in which rats waited for partners increased in both success trials and failure trials (Supplementary Figure S2E). In the 1sCo phase, the duration of social time window showed a weak positive correlation with SR (Supplementary Figure S2F). We also analyzed the head direction (α angle) and social orientation (β angle; see Methods and reference ^64^; Figure 2B, bottom) during these time windows. Both the head direction and the social orientation decreased across training (Supplementary Figure S2G). Notably, in the 1sCo phase, the social orientation showed a moderate negative correlation with SR (r = −0.64; Supplementary Figure S2H). These results suggest that rats actively awaited partners and increased attention to them within the nose-poke area.

To quantify social interactions during these time windows, we analyzed dyadic nose approach behavior, defined as when rats’ nose passed through the perforated partition (Supplementary Figure S2I). During cooperative training, the cumulative duration of nose approach increased progressively in both success trials and failure trials (1sCo D12: 1.87 s and 1.60 s, respectively; Figure 2D), and showed a weak positive correlation with SR (Supplementary Figure S2J). The proportions of both trial types involving nose approach also increased gradually (Supplementary Figure S2K). Moreover, the proportion of time spent in nose approach increased in both the waiting and social time windows during the 1sCo phase (Supplementary Figure S2L). Notably, the probability of nose approach was significantly higher on 1sCo D6 and D12 than on D1, with its peak occurring in the middle of the social time window (Figure 2E).

To further quantify social interactions between dyads, we installed infrared detectors in the middle of cages (Supplementary Figure S2M) to record partial physical contacts close to the nose-poke port (e.g., forepaw or nose). Both the total interaction bouts and the mean interaction duration increased during cooperative training (Supplementary Figure S2N). Moreover, the proportion of success trials involving social interactions gradually increased to nearly 70% (Supplementary Figure S2O), with most interactions occurring within 2 s before successful cooperation (Supplementary Figure S2P). To examine the necessity of these social interactions for efficient cooperation, metal meshes were installed in the middle of cages, specifically blocking physical contacts within the NPA. We found that blocking social interactions by addition of the mesh significantly reduced SR (67.06% to 22.84%; Figure 2F) and increased NPI (337.21 ms to 434.59 ms; Figure 2G) in well-trained dyads. Subsequent removal of the mesh restored both SR and NPI to well-trained levels (Figure 2F, G). Collectively, these results demonstrate that rats actively await partners and engage in social interactions to achieve efficient cooperation.

### Rats primarily cooperate through communication-based strategy

Our observations indicate that the cooperative processes of different dyads are diverse, suggesting that they may utilize multiple strategies to achieve cooperation. To investigate the formation of cooperative strategies in rats, we classified their success trials based on the durations of waiting time window (WTW) and social time window (STW), and the interval between leaving the water lick area (ΔWLA; see Methods). Three distinct cooperative strategies were identified: the communication strategy (Type 1; STW>1.5 s), the short social strategy (Type 2; characterized by STW≤1.5 s, and either WTW or ΔWLA>1.5 s), and the synchrony strategy (Type 3; defined by all of ΔWLA, WTW, and STW≤1.5 s; Figure 3A–C; Supplementary Figure S3A–D). By analyzing behavioral parameters across cooperative strategies, we found that the communication strategy showed a significantly lower NPI than the synchrony strategy in the late 1sCo stage (Supplementary Figure S3E). Compared to the synchrony strategy, both the communication and short social strategies exhibited longer intervals between leaving the WLA or entering the NPA (namely waiting time window) and more frequent nose approaches (Supplementary Figure S3F–H). Notably, the communication strategy had a significantly longer social time window than the other two strategies (1sCo D12: 2.66 s; Supplementary Figure S3I).

**Figure 3.**
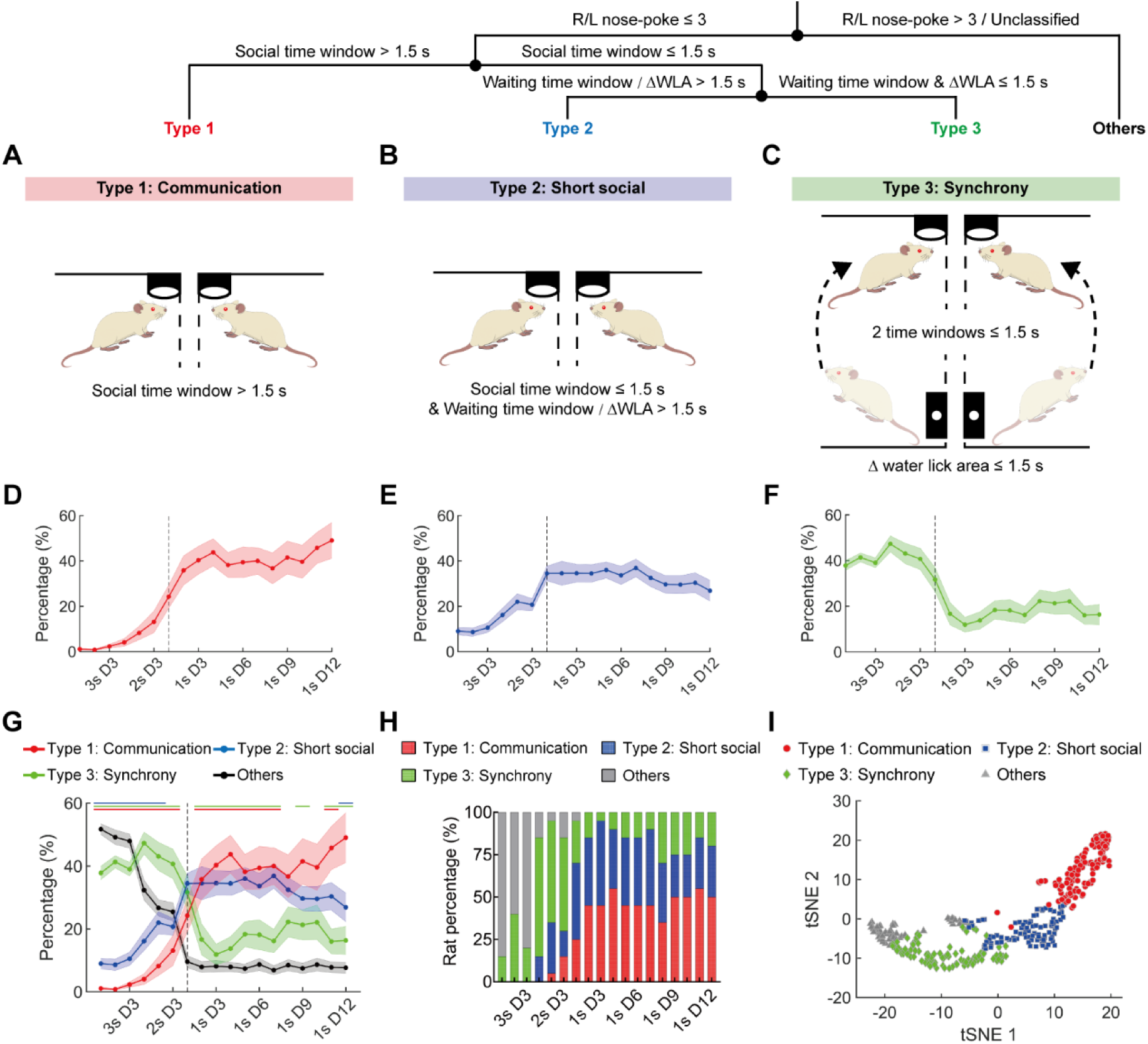
Classification of cooperative strategies during cooperative training. A–C: Schematics of three different types of cooperative strategies: communication (Type 1, A), short social (Type 2, B), and synchrony (Type 3, C). Top panel shows the classification process of cooperative strategies. D–F: Percentage changes in three cooperative strategies during cooperative training (*n*=20; D, Type 1; E, Type 2; F, Type 3). G: Percentage changes in different cooperative strategies during cooperative training (*n*=20). Lines at the top indicate statistical significance (*P*<0.05): Type 1 vs. Type 2 (blue), Type 1 vs. Type 3 (green), and Type 2 vs. Type 3 (red). H: Percentage of dyads using the dominant strategies during cooperative training. I: t-SNE plot of all sessions with different dominant strategies during cooperative training (all sessions, *n*=360). In D–G, vertical dashed lines indicate D1 of 1sCo. Data are presented as mean ± SEM. Statistics: one-way ANOVA (G).

The proportional changes in each strategy throughout cooperative training were quantified (Figure 3D–G). The proportion of the communication strategy increased progressively (1.11% to 49.05%; Figure 3D, G) and showed a weak positive correlation with SR (Supplementary Figure S3J). In contrast, the short social strategy proportion initially increased but subsequently declined (9.02%, 34.50% to 26.89%; Figure 3E, G), showing a weak negative correlation with SR (Supplementary Figure S3K). While predominant during the 3sCo and 2sCo phases, the synchrony strategy proportion rapidly declined upon entering the 1sCo phase (31.63% to 16.36%; Figure 3F, G), exhibiting a weak positive correlation with SR (Supplementary Figure S3L). Notably, the combined proportion of communication and synchrony strategies revealed a moderate positive correlation with SR during the 1sCo phase (r = 0.38; Supplementary Figure S3M). We also calculated the strategy transition probability from the present to the subsequent success trial, and found that the probability of communication to communication gradually increased during the 1sCo phase (Supplementary Figure S3N–P).

We further defined the strategy with the highest proportion as the dominant strategy for each dyad. Analysis of dominant strategies revealed the proportions of dyad dominated by the communication, short social, and synchrony strategies were 50%, 30%, and 20% respectively, on day 12 of the 1sCo phase (Figure 3H). Moreover, visualization of dominant strategies by t-distributed stochastic neighbor embedding (t-SNE) showed that sessions dominated by these strategies formed three continuous yet distinct clusters (Figure 3I). We also set the cooperative-strategy cutoffs to 1 s and 2 s, respectively, and found that a 1-s cutoff increased the proportion of the communication strategy during the 1sCo stage, making it consistently the most predominant strategy (Supplementary Figure S3Q, R). In contrast, a 2-s cutoff caused the three strategies to show similar proportions in the late 1sCo stage, although the proportion of dyad dominated by the communication strategy still reached 50% at 1sCo D12 (Supplementary Figure S3S, T). Taken together, these results demonstrate that rats utilize multiple cooperative strategies to achieve cooperation, with the communication strategy emerging as the predominant strategy over time.

Importantly, compared with our previous 3s-1s task (7 days of 3sCo and 14 days of 1sCo) ^25^, our improved 3s-2s-1s task includes a 20-cm divider in the center of each chamber (Figure 1A), which reduces synchronous movement between rats (Supplementary Figure S4). Joint analyses (Supplementary Figure S4A–C) showed that SR of the 3s-1s task did not differ from that of the 3s-2s-1s task (Supplementary Figure S4B). However, unlike in the 3s-2s-1s task (Figure 2C), the waiting time window and social time window in the 3s-1s task did not increase over training (Supplementary Figure S4D). Most notably, whether using a 1.5-s or 1-s cutoff (a stricter criterion for synchrony), the synchrony strategy in the 3s-1s task remained significantly higher than the other two cooperative strategies (Supplementary Figure S4E–H), in clear contrast to our improved 3s-2s-1s task (Figure 3G, H; Supplementary Figure S4I–K).

**Figure 4.**
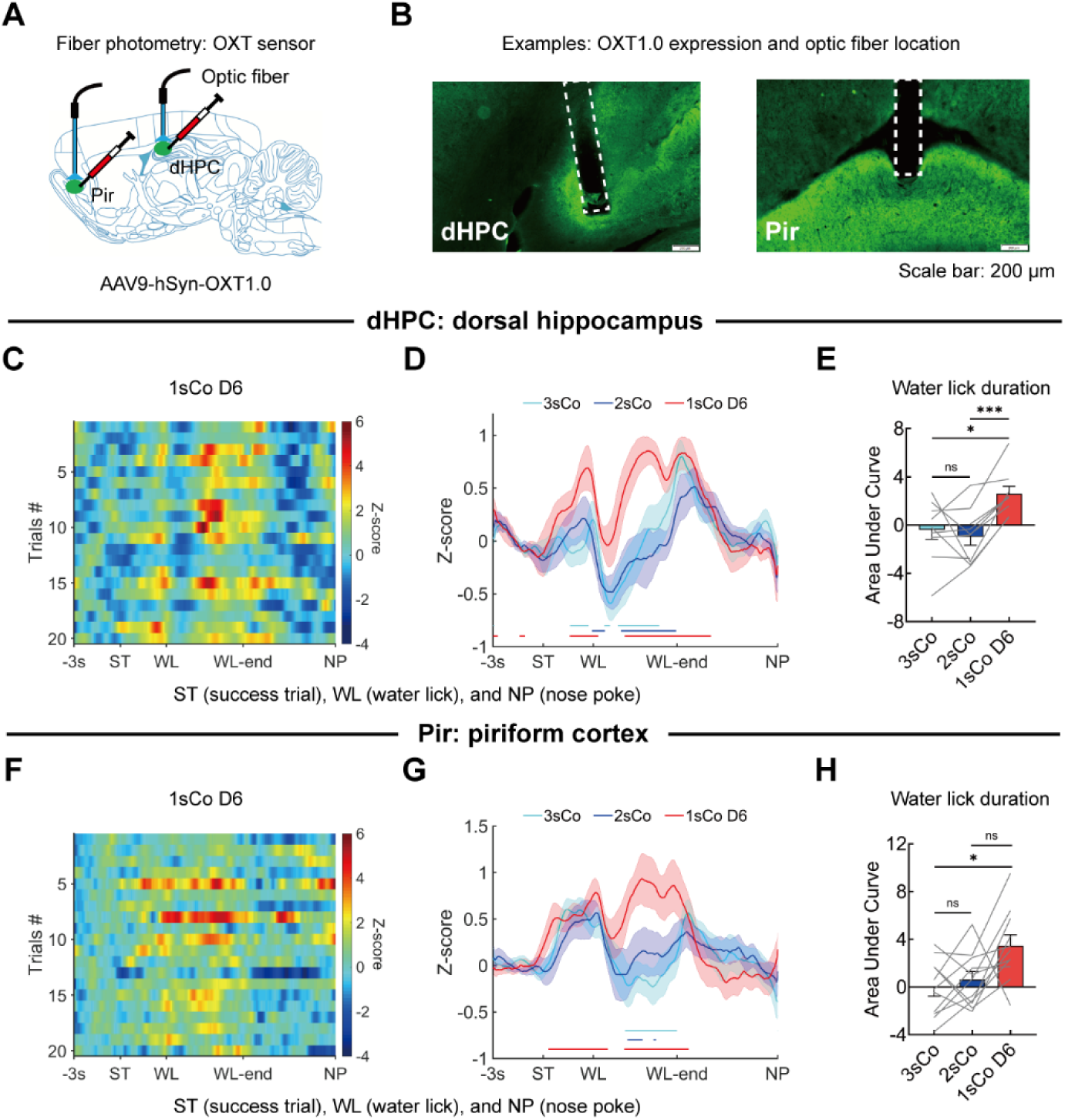
Release dynamics of OXT in dHPC and Pir during cooperative training. A: Unilateral OXT1.0 expression in the Pir and dHPC for fiber photometry recording. B: Representative OXT1.0 expression and optic fiber location in the dHPC and Pir. Scale bar, 200 μm. C: Heatmap of OXT signals in the dHPC during D6 of 1sCo (*n*=20, from one rat). D: OXT signal dynamics in the dHPC across three learning stages of cooperation (3sCo, 2sCo, and 1sCo D6; *n*=9–10). E: Among three learning stages, comparison of OXT signal differences in the dHPC at water lick duration (*n*=9–10; 3sCo vs. 2sCo, *P*=0.7990; 3sCo vs. 1sCo D6, *P*=0.0187; 2sCo vs. 1sCo D6, *P*=0.0005). F: Heatmap of OXT signals in the Pir during D6 of 1sCo (*n*=20, from one rat). G: OXT signal dynamics in the Pir across three learning stages of cooperation (3sCo, 2sCo, and 1sCo D6; *n*=11). H: Among three learning stages, comparison of OXT signal differences in the Pir at water lick duration (*n*=11; 3sCo vs. 2sCo, *P*=0.7380; 3sCo vs. 1sCo D6, *P*=0.0209; 2sCo vs. 1sCo D6, *P*=0.0954). In D and G, lines at the bottom indicate statistical significance (*P*<0.05): 1sCo D6 vs. baseline (red), 1sCo D6 vs. 3sCo (cyan), and 1sCo D6 vs. 2sCo (blue). Abbreviation: ST (success trial), WL (water lick) and NP (nose poke). Data are presented as mean ± SEM. Statistics: one-way ANOVA (D and G) and repeated measures one-way ANOVA (E and H). ns: Not significant; *: *P*<0.05; **: *P*<0.01; ***: *P*<0.001.

### Oxytocin release is increased along with the acquisition of cooperation

Previous studies have shown that OXT modulates social recognition and learning via the dorsal hippocampus (dHPC) and piriform cortex (Pir) in mice ^65–67^. To determine whether OXT contributes to cooperative behavior in rats, we measured OXT release in the dHPC and Pir during cooperative training using a genetically encoded OXT sensor (GRAB_OT1.0_ or OXT1.0) ^68^. In naïve rats, we unilaterally injected the AAV9-hSyn-OXT1.0 virus into the dHPC and Pir (Figure 4A, B) and recorded OXT signal from these regions at three training stages: 3sCo, 2sCo, and the mid-phase of 1sCo (day 6 of 1sCo, 1sCo-D6).

During the 1sCo-D6, OXT release in the dHPC increased following successful cooperation (water lick latency, from success trial to water lick, ST-WL), decreased and then rebounded during water licking (water lick duration, WLD; from WL to WL-end), and finally declined after licking ended (Figure 4C, D). Notably, during the learning process of cooperation, OXT release in the dHPC was significantly elevated in 1sCo-D6 compared to 3sCo or 2sCo (Figure 4D), with a specific enhancement during the WLD period (Figure 4D, E). The OXT elevation in the 1sCo phase indicates that OXT release is not merely resulted from water reward (3sCo and 2sCo phases have the same reward), but may be correlated with task difficulty. Similar dynamics of OXT release were observed in the Pir (Figure 4F–H). Collectively, these results indicate that the dynamics of OXT release are correlated with the acquisition of cooperation.

### Oxytocin deficiency impairs rats’ acquisition of cooperation

To further determine whether OXT is required for efficient cooperation, we trained *Oxt* knockout rats (OXT^⁻/⁻^, OXT-KO) ^69,70^ and their wild-type littermates (OXT⁺/⁺, WT-LM) in the cooperative task. Knockout efficiency was verified by immunohistochemistry (Figure 5A). We first assessed the basic behavioral phenotypes of adult OXT-KO rats. In the open-field test (OFT), locomotor distance was comparable between OXT-KO rats and WT-LM (Figure 5B). Similarly, time spent in the open arms of the elevated plus maze (EPM) and in the center of the OFT did not differ between groups (Supplementary Figure S5A, B). These results suggest that OXT-KO rats displayed normal locomotion and anxiety-like behavior. In the three-chamber test (TCT), OXT-KO rats exhibited normal social preference (conspecifics versus toy) compared to WT-LM (Figure 5C; Supplementary Figure S5C). However, unlike WT-LM, OXT-KO rats failed to discriminate between familiar and novel conspecifics in the TCT (Figure 5D; Supplementary Figure S5D), indicating impaired social novelty recognition.

**Figure 5.**
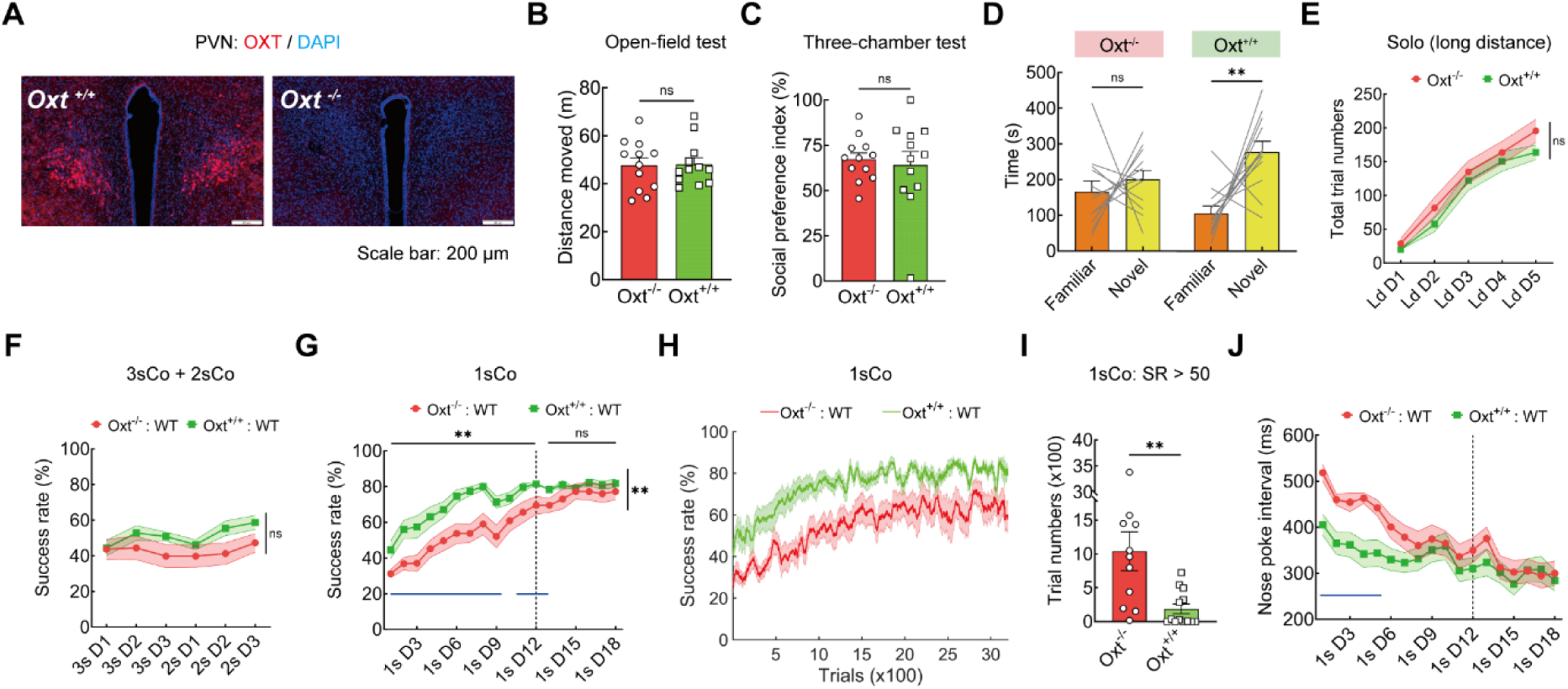
Behavioral phenotype of OXT knockout rats during social tests and cooperative training. A: OXT protein expression in the PVN of WT littermate (WT-LM, OXT^+/+^) and OXT-KO (OXT^-/-^) rat. Scale bar, 200 μm. B–C: In OXT-KO (red, *n*=12) and WT-LM (green, *n*=12) rats, total distance moved during open-field test (B, *P*=0.9227) and social preference index during three-chamber test (TCT, C, *P*=0.7080). D: Time spent in familiar versus novel sides of OXT-KO and WT-LM rats during TCT (*n*=12 and 12; OXT-KO, *P*=0.5352; WT-LM, *P*=0.0051). E: Trial numbers of OXT-KO and WT-LM rats during Solo training (*n*=13 and 13, *P*=0.2559). F: Success rate (SR) of OXT-KO and WT-LM dyads during 3sCo and 2sCo training (*n*=13 and 13, *P*=0.1533). G: SR of OXT-KO and WT-LM dyads during 1sCo training (*n*=11 and 13; 1s D1–18, *P*=0.0042; 1s D1–12, *P*=0.0014; 1s D13–18, *P*=0.2474). H–I: SR of OXT-KO and WT-LM dyads (H, calculated by trials) and trials required to achieve >50% SR (I, *P*=0.0019) during 1sCo training. J: Nose-poke interval of OXT-KO and WT-LM dyads during 1sCo training (*n*=11 and 13). In G and J, vertical dashed lines indicate D12 of 1sCo. In G and J, blue lines at the bottom indicate statistical significance (*P*<0.05). Data are presented as mean ± SEM. Statistics: unpaired *t*-tests (B, G and J), unpaired *t*-test with Welch’s correction (C), Mann Whitney test (I), paired *t*-tests (D), and two-way ANOVA (E–G). All statistical tests are two-tailed (except for two-way ANOVA). ns: Not significant; *: *P*<0.05; **: *P*<0.01; ***: *P*<0.001.

For cooperative training, OXT-KO rats or WT-LM were paired with naïve SD partners. We found that the learning curves of both dyads were similar during pre-cooperation training (solo; Figure 5E) and the 3sCo-2sCo phases (Figure 5F). However, during the 1sCo phase, the OXT-KO dyads showed significantly slower improvement in success rate (SR; Figure 5G; red vs. green: 1s D1–18, *P*=0.0042; 1s D1–12, *P*=0.0014) and required more trials to achieve an SR exceeding 50% (Figure 5H, I). Further analysis revealed that while the numbers of success trials were similar (Supplementary Figure S5E), OXT-KO dyads exhibited significantly more failure trials during the first 14 days of 1sCo (Supplementary Figure S5F). However, the ratios of each rat initiating nose-poking were equal for both success and failure trial in OXT-KO or WT-LM dyads (Supplementary Figure S5G, H). During the first 5 days of 1sCo, the OXT-KO dyads showed significantly longer NPI than WT-LM dyads (red vs. green; Figure 5J). Nevertheless, both SR and NPI became comparable between dyads later in the 1sCo phase (SR: 1s D13–18, *P*=0.2474; Figure 5G, J). Together, these results demonstrate that OXT deficiency impairs the acquisition of cooperation in rats.

### Oxytocin system is required for the formation of communication-based strategy

To determine whether OXT mediates the formation of cooperative strategies, we performed joint analyses of behavioral events and video tracking from OXT-KO and WT-LM dyads during the 1sCo phase. During the late stage of training, the OXT-KO dyads showed significantly shorter social time window and waiting time window than the WT-LM dyads (red vs. green; Figure 6A; Supplementary Figure S6A), indicating less social interaction. Strategy analysis revealed that OXT knockout shifted strategy adoption from a predominant reliance on communication strategy in the WT-LM dyads to a more balanced adoption of all three strategies in the OXT-KO dyads (Figure 6B; Supplementary Figure S6B). On day 18 of 1sCo, the OXT-KO dyads used communication strategy less and synchrony strategy more than the WT-LM dyads (red vs. green; Figure 6C; Supplementary Figure S6C). We also set the cooperative-strategy cutoffs to 1 s and 2 s, and found that changing the cutoff did not affect the above result: the OXT-KO dyads still used the communication strategy less than the WT-LM dyads (Supplementary Figure S6D–G). Together, these results indicate that OXT mediates the formation of cooperative strategies, and its deficiency impairs the adoption of the communication strategy.

**Figure 6.**
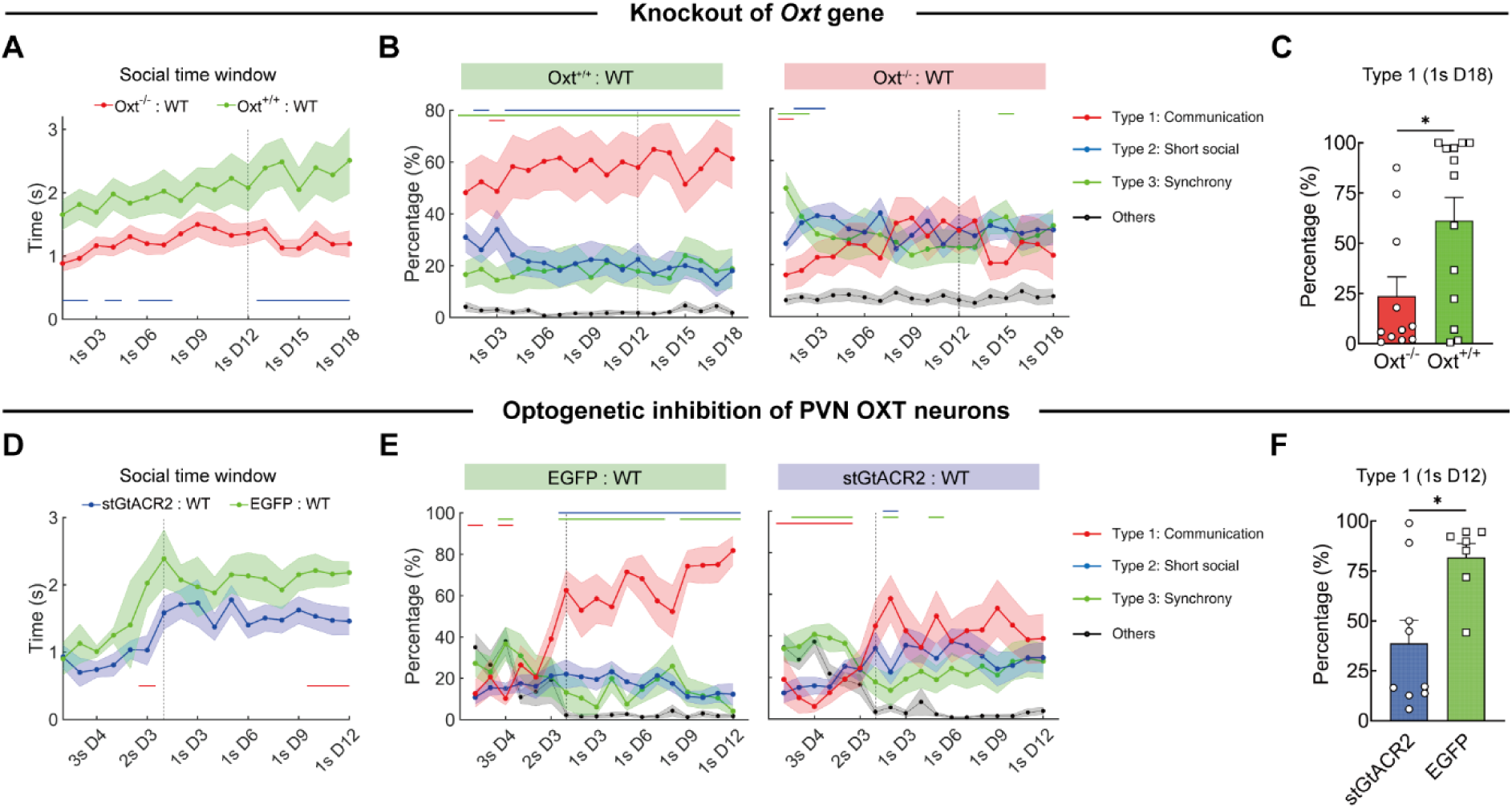
OXT system mediates the formation of cooperative strategies during learning. A: Durations of the social time window in OXT-KO and their WT littermate (WT-LM) dyads during 1sCo training (*n*=11 and 13). B: Percentage changes of three cooperative strategies in WT-LM (left, *n*=13) and OXT-KO dyads (right, *n*=11) during 1sCo training. C: Percentage of strategy Type 1 (communication) in OXT-KO and WT-LM dyads on D18 of 1sCo (*n*=11 and 13, *P*=0.0351). D: Durations of the social time window in stGtACR2 and control dyads during cooperative training (*n*=9 and 7). E: Percentage changes of three cooperative strategies in control dyads (left, *n*=7) and stGtACR2 dyads (right, *n*=9) during cooperative training. F: Percentage of strategy Type 1 (communication) in stGtACR2 and control dyads on D12 of 1sCo (*n*=9 and 7, *P*=0.0418). In A and B, vertical dashed lines indicate D12 of 1sCo. In D and E, vertical dashed lines indicate D1 of 1sCo. In A and D, blue/red lines at the bottom indicate statistical significance (*P*<0.05). In B and E, lines at the top indicate statistical significance (*P*<0.05): Type 1 vs. Type 2 (blue), Type 1 vs. Type 3 (green), and Type 2 vs. Type 3 (red). Data are presented as mean ± SEM. Statistics: one-way ANOVA (A–B and D–E) and two-tailed Mann Whitney test (C and F). *: *P*<0.05.

The PVN OXT-ergic neurons are a primary source of OXT and project to the dHPC and Pir ^71–73^. To determine whether PVN OXT-ergic neurons also mediate the strategy formation, we first examine their contribution to OXT release within the dHPC and Pir, and then continuously inhibit them during cooperative training. Synaptophysin-EGFP was expressed in PVN OXT-ergic neurons of OXT-Cre rats (Supplementary Figure S7A), and EGFP signals were detected in both dHPC and Pir (Supplementary Figure S7B), indicating that PVN OXT-ergic neurons project to these regions. We further optogenetically activated or inhibited PVN OXT-ergic neurons while simultaneously recording OXT sensor signals in the dHPC and Pir (Supplementary Figure S7C). Optogenetic activation significantly elevated OXT release in both dHPC and Pir, whereas inhibition suppressed it (Supplementary Figure S7D, E). These results confirm that PVN OXT-ergic neurons contribute to OXT release in the dHPC and Pir.

Next, to continuously inhibit PVN OXT-ergic neurons during cooperative training, we bilaterally injected AAV9-hSyn-DIO-stGtACR2 or AAV9-hSyn-DIO-EGFP (control) virus into the PVN of OXT-Cre rats (Supplementary Figure S8A). DIO-EGFP expression specificity and stGtACR2 efficacy were validated (Supplementary Figure S8B, C). For cooperative training, stGtACR2 rats or control rats were paired with naïve SD partners. Optogenetic inhibition was applied during the water lick latency and water lick duration (WLL-WLD; Supplementary Figure S8D), as OXT release increases within this time window (Figure 4D, G). During the late stage of 1sCo training, inhibition of PVN OXT-ergic neurons significantly reduced the social time window (blue vs. green; Figure 6D), without affecting the waiting time window (Supplementary Figure S8E). Furthermore, inhibition of PVN OXT-ergic neurons shifted strategy adoption from a predominant reliance on communication strategy in control dyads to a more balanced adoption of all three strategies in stGtACR2 dyads (Figure 6E; Supplementary Figure S8F). On day 12 of 1sCo, stGtACR2 dyads used communication strategy less and synchrony strategy more than control dyads (blue vs. green; Figure 6F; Supplementary Figure S8G). We also set the cooperative-strategy cutoffs to 1 s and 2 s, and found that under the 2-s cutoff condition, stGtACR2 dyads still used the communication strategy less than control dyads (Supplementary Figure S8H–K). Collectively, these results demonstrate that PVN OXT-ergic neurons also mediate the formation of cooperative strategies, and their continuous inhibition impairs the adoption of the communication strategy.

## DISCUSSION

To investigate the learning process and strategy formation underlying rat cooperation, as well as the regulatory role of OXT in this behavior, we established a temporal coordination-based cooperative task and manipulated the OXT system during training. Through progressive cooperative training, rats acquired cooperation and efficiently performed the task. Furthermore, rats waited for their partners and engaged in social interactions, which were essential for efficient cooperation performance. During the cooperative training, rats displayed different cooperative strategies, with the communication-based strategy gradually increasing and eventually becoming predominant. The cooperative learning process was accompanied by elevated OXT release in brain regions associated with social learning (dHPC and Pir), whereas OXT knockout impaired learning speed. Moreover, both OXT knockout and continuous inhibition of PVN OXT-ergic neurons during learning reduced the adoption of the communication-based strategy.

### Rats’ behavioral performance during cooperative training

To investigate the learning process and strategy formation in rat cooperation, we combined behavioral events with video tracking to quantify the dynamics of temporal coordination (Figure 1), social interactions (Figure 2), and cooperative strategies (Figure 3). Our results showed that, through progressive cooperative training, paired rats achieved high task performance (Figure 1 and 7, left), comparable to previous work under similar conditions (1-second time window) ^25,31^. Notably, better temporal coordination (reflected by a shorter nose-poke interval) positively correlated with higher success rates (SR). During training, rats exhibited an increasing waiting time window while reducing head orientation (toward the central perforated panel), indicating increased attention to the partner’s position. Such waiting behavior has also been observed in cooperative tasks involving mice and elephants ^16,23^, and is considered evidence that animals recognize the need for a partner ^16^. We further observed that both the social time window and nose approach behavior increased over training, while social orientation decreased. Nose approach behavior showed a positive correlation with SR, reinforcing findings from previous rodent studies ^23,25^. When physical contact was prevented by installing a metal mesh, cooperation efficiency significantly decreased, confirming the importance of physical contact ^28,74^. However, even without physical contact, SR and NPI remained significantly different from those in shuffle controls, suggesting that other sensory modalities (e.g., vision, olfaction) also contribute to cooperation ^27^.

**Figure 7.**
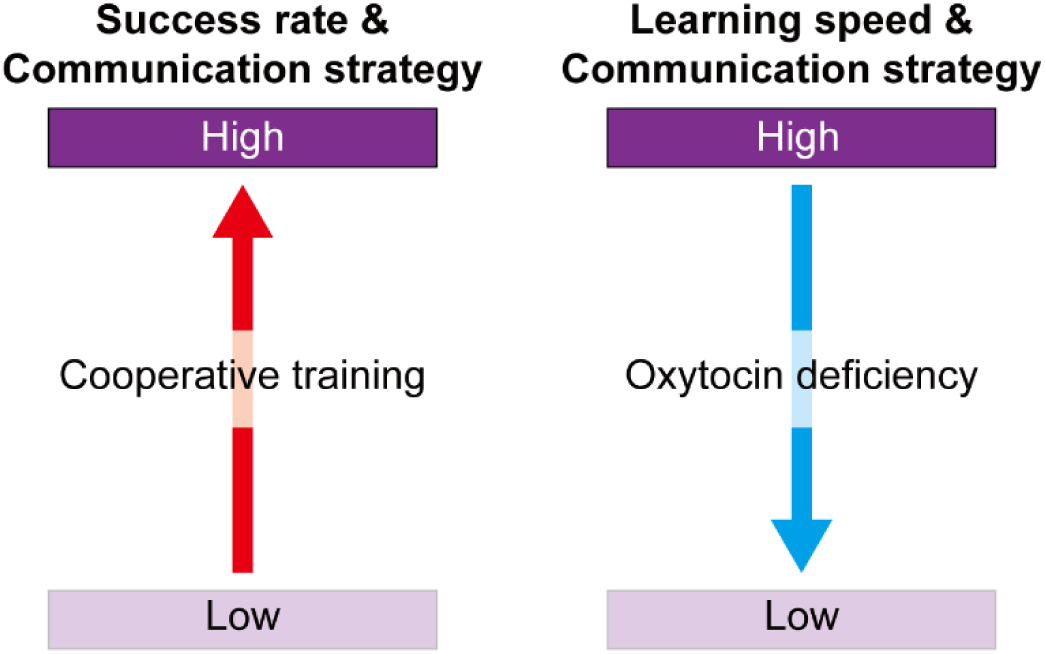
Summary of the functions of OXT during cooperative training. Left: Progressive cooperative training improves the cooperative abilities of rats, as indicated by an increased success rate and greater adoption of the communication-based cooperative strategy. Right: Oxytocin deficiency impairs the cooperative abilities of rats, as indicated by reduced learning speed and reduced adoption of the communication-based cooperative strategy.

### Cooperative strategies formation and cooperation level of rats

We identified three distinct cooperative strategies in rats: communication, short social, and synchrony (Figure 3). In the 1sCo phase, the communication and synchrony strategies (individually or combined) were positively correlated with SR. During training, the synchrony strategy was predominant in the 3sCo-2sCo phase but declined rapidly to around 20% after entering the 1sCo phase. In contrast, the communication strategy gradually increased to around 50%, becoming the predominant strategy (Figure 3 and 7, left). Previous studies categorizes chimpanzee (*Pan troglodytes*) cooperation into four progressively complex levels: similarity, synchrony, coordination, and collaboration ^1^; or alternatively, by-product collaboration, socially influenced collaboration, actively coordinated collaboration, and collaboration based on shared intentionality ^5^. Behavioral evidence from chimpanzees and elephants, including waiting for partners and recognizing when a partner can perform its role, suggests that their cooperation can reach the level of actively coordinated collaboration ^5^. Based on our findings, we propose that the synchrony strategy observed in rats corresponds to the second level of cooperative hunting (synchrony), while the communication strategy may correspond to the third level (coordination or actively coordinated collaboration).

Importantly, in our previous 3s-1s task without a middle divider (21 days) ^25^, paired rats preferred completing the cooperative task through synchronous movements (level 2); whereas in the improved 3s-2s-1s task with an added middle divider (18 days), paired rats showed a stronger preference for completing the task via social communication (level 3). Therefore, to some extent, our improved 3s-2s-1s task enhances the level of cooperation in paired rats. Recently, different strategy analyze has been applied to cooperative behavior (joint lever-pulling) in common marmosets (*Callithrix jacchus*), which is highly analogous to the nose-poke task used here ^75^. During the joint lever-pulling task, marmosets exhibit strategies such as gaze-dependent coordination (gaze-and-pull strategy) and gaze-independent rhythmic pulling (pull-in-rhythm strategy) ^75^, which could be conceptually linked to the communication and synchrony strategies observed in rats. Taken together, these studies extend current understanding in the field and provide new experimental evidence for exploring the evolution of cooperation through cross-species comparisons.

### The role of oxytocin in the acquisition of cooperation

To examine whether the learning process of cooperation in rats is regulated by OXT, we recorded and manipulated the OXT system during cooperative training. Our results revealed that OXT release dynamics correlate not only with task execution but also with the acquisition of cooperation (Figure 4). OXT knockout (OXT-KO) rats exhibited deficits in social novelty recognition and cooperative learning (Figure 5 and 7, right). Previous studies using OXT sensors have demonstrated that OXT release is elevated during courtship behavior ^68^ and exhibits temporal fluctuations (OXT oscillations) ^76^. In the prelimbic cortex (PrL), OXT release progressively increases during transitions from non-REM sleep to REM (rapid-eye-movement) sleep and then to wakefulness, contributing to social recognition ^77^. However, previous studies in animals only showed that urinary OXT levels are elevated after cooperative behaviors ^56–58^. Therefore, our results further reveal the temporal dynamics of OXT release during the acquisition and execution of cooperative behavior.

The social novelty deficits observed in our OXT-KO rats (Figure 5) are consistent with earlier findings in OXT-KO mice ^41^. In a prosocial behavior paradigm, blocking oxytocin receptors (OXTR) in the anterior cingulate cortex (ACC) impaired the acquisition of helping behavior in rats ^78^. So far, prior findings indicate OXT is essential for social learning (or recognition) ^79^, that this role is evolutionarily conserved ^80,81^ and mediated by multiple brain regions, including the PrL ^77,82^, Pir ^65^, and HPC ^66,67^. Therefore, our results extend OXT’s role from basic social recognition to the acquisition of complex social behaviors.

### The role of oxytocin in the formation of cooperative strategies

During cooperative learning, both OXT knockout and continuous inhibition of PVN OXT-ergic neurons reduced the adoption of the communication strategy (Figure 6 and 7, right), resulting in a more balanced use of all three strategies. In wild-type rats, both communication and synchrony strategies (individually or combined) were positively correlated with SR. Thus, manipulating the OXT system altered strategy choice during cooperative learning (reducing communication while increasing synchrony and short social strategies), without affecting SR once the task was well learned (Figure 5). To dissect the underlying social-cognitive mechanisms, we propose that strategy formation in paired rats may be mediated by social reward. Previous studies have shown that OXT is necessary for social conditioned place preference (CPP) in mice ^83^, and that optogenetic or chemogenetic activation of PVN OXT-ergic neurons enhances social preference (or reward) toward conspecifics ^43,44,65,84^. Therefore, OXT knockout or inhibition of PVN OXT-ergic neurons may reduce social reward, thereby decreasing preference for the communication strategy. Although our male OXT-KO rats exhibited normal social preference, earlier work has reported that while male OXT-KO mice also show normal social preference ^85^, male OXTR-KO mice exhibit social preference deficits ^86,87^. Moreover, the three-chamber social test used in these studies measures short-term social preference, whereas cooperation can be influenced by longer-term social relationships ^88,89^. We therefore propose that OXT may affect cooperative strategy formation not only through modulation of social reward but also by shaping social relationships (e.g., peer bonding ^70,90^).

In summary, our results demonstrate that OXT mediates both the learning process and the strategy formation underlying cooperative behavior in rats, thereby enabling them to rapidly acquire cooperation and obtain mutual rewards. Children with autism spectrum disorder (ASD) often exhibit social deficits and impaired cooperative abilities ^91,92^. Given that OXT has therapeutic potential for ASD ^93,94^, supplemental OXT administration during social task learning ^95^ may facilitate skill acquisition and ameliorate autism symptoms in affected children.

### Limitations of the study

In terms of behavior, the extent to which rats recognize their partners’ roles remains unclear. This question is closely related to whether they develop shared intentionality ^96^. Regarding neural mechanisms, because all experiments were conducted in male rats and OXT exhibits sexual dimorphism ^97,98^, it remains to be determined whether OXT has similar effects in females. Furthermore, we manipulated only PVN OXT-ergic neurons, whereas OXT-ergic neurons in the SON may also contribute to the learning of cooperative behavior ^99^. Likewise, whether downstream PVN circuits expressing OXTR are involved in cooperative learning remains unknown. Although we observed enhanced OXT release in the dHPC and Pir, causal manipulations are needed to establish the functional contributions of these regions during learning. Finally, the effects of OXT on social cognition and behavior are strongly influenced by contextual factors and individual differences ^100,101^. While our study focused on OXT’s role in affiliative cooperation, it provides a foundation for future research into how OXT function may differ between affiliative and antagonistic contexts, and under conditions of positive versus negative valence ^102,103^.

## METHODS

### Animals

Male wild-type Sprague-Dawley (SD) rats were purchased from Shanghai SLAC Laboratory Animal Co., Ltd. Oxytocin knockout (OXT-KO) SD rats, generated via CRISPR-mediated deletion ^69,70^ were provided by Prof. Pu Fan (Peking Union Medical College). OXT-Cre transgenic SD rats, in which a P2A-iCre cassette was inserted upstream of the 3’ untranslated region (UTR) of the *Oxt* gene ^84^, were generated by Beijing Biocytogen Co., Ltd. Rats were maintained under a standard 12-h light/dark cycle (lights on at 07:00). All animals were used and operated in accordance with the regulations of the Institutional Animal Care and Use Committee of the Center for Excellence in Brain Science and Intelligent Technology (Institute of Neuroscience), Chinese Academy of Sciences.

For cooperative training, rats were randomly paired and housed together. Non-surgical rats began training at 6 weeks of age (180–200 g), whereas surgical rats began at 9 weeks of age (300–350g). All animals were water-deprived for >20 h before training. On training days, each rat received 3–5 min of water access within 1–2 h after the training session (sufficient to maintain weight gain). On non-training days, water was provided *ad libitum*.

### Development of the cooperative training system

The cooperative training system consisted of a behavioral apparatus and a control unit (Anilab Software & Instruments, China). Two behavioral cages (40×30×45 cm, L×W×H) were placed 1.5 cm apart within a sound-attenuating, ventilated cabinet. Each cage was constructed from five panels and equipped with a nose-poke port and a water port, both with infrared detectors linked to the AniLab controller (input). The adjoining panels were perforated in a rectangular pattern (1.5×15 cm, W×H; gap 0.5 cm) and fitted with a 20-cm divider to partition each cage into inner and outer compartments. Water ports were connected to pumps via tubing. Two pumps, two LEDs, and a buzzer were linked to the AniLab controller (output). The cabinet also housed top- and side-mounted cameras and a speaker, connected to the control computer. The computer was connected to the AniLab controller via USB and ran training protocols (solo or cooperative) programmed in LabState software (Anilab Software & Instruments, China). At the start of each session, paired rats (dyads) were placed in their assigned cages, and the task was started. Both cages were illuminated (~10 lux), and infrared detectors, pumps, cameras, and sound devices were activated. Sessions lasted 30 min, after which lights and recording stopped automatically.

To measure physical contacts between dyads, an additional infrared detector was installed on the outer side of the perforated panel, slightly below the nose-poke height, to detect contacts within 3 cm along the perforated panel. To block tactile but not visual, olfactory, or auditory interaction, metal meshes (0.9-mm aperture) were installed over the perforated panels.

### Training protocol of rat cooperation task

The training procedure consisted of two stages: pre-cooperation training (Stage 1) and progressive cooperative training (Stage 2). The cooperative task was based on temporal coordination, requiring two paired rats to simultaneously nose-poke within a defined time window to obtain rewards. Depending on the duration of time window, the task was classified as 3-second cooperation (3sCo), 2-second cooperation (2sCo), and 1-second cooperation (1sCo).

Stage 1 (pre-cooperation training): This stage included short-distance training (water port is close to the nose-poke port) and long-distance training (water port positioned farther away). Rats were trained to nose-poke individually for water rewards (0.5‰ saccharin sodium salt hydrate, Sigma-Aldrich, Germany). When the infrared detector in the nose-poke port was triggered, a pump delivered a drop of water (~20 μL) accompanied by an auditory cue (right cage, 5-kHz tone; left cage: 1-kHz tone). After a rat entered the water port, the trial was counted, and the program reset. In short-distance training (~4 days), rats were considered well-trained once they completed >100 trials in a single session. In long-distance training (~5 days), the water port was placed on the opposite side of the nose-poke port (inner versus outer side of the cage; see Figure 1A). Rats were considered well-trained when they completed >50 trials in three consecutive sessions, after which progressive cooperation training began.

Stage 2 (progressive cooperative training): This stage consisted of 3 days of 3sCo, 3 days of 2sCo, and 12–18 days of 1sCo. Rats were trained to simultaneously nose-poke within the defined window (3 s, 2 s, or 1 s) with their partner to receive water rewards. In 3sCo, if one rat nose-poked and its partner did so within 3 s, the trial was counted as successful (success trial, ST), triggering a 1-s auditory cue (8-kHz tone) and one water drop for each rat. If the partner failed to nose-poke within 3 s, the trial was counted as a failure (failure trial, FT) and a 1-s white noise (~70 dB) was played. After the auditory cue, a 1-s inter-trial interval (ITI) was applied before the program reset. The same contingencies applied for 2sCo and 1sCo, with the time window adjusted to 2 s and 1 s, respectively.

Extended 3sCo training (18 days) included two phases: pre-cooperation training followed by 18 consecutive days of 3sCo, identical in procedure to the 3sCo phase within progressive cooperation training. In manipulation experiments on cooperative learning (including OXT-KO rats and optogenetic inhibition of PVN OXT-ergic neurons), experimental and control rats were evenly distributed across behavioral cages. Dyads completing fewer than ~10 trials in most 3sCo-2sCo sessions did not proceed to 1sCo training.

### Analysis of parameters related to cooperative behavior

The success rate (SR) was calculated as: Success rate=success trials/(success trials+failure trials)×100%. The initiator ratio of success trial (ST) or failure trial (FT) was calculated as: Initiator ratio=R/(R+L), where R denotes trials initiated by the right side, and L denotes trials initiated by the left side. The nose-poke latency (NPL) was defined as the interval between the end of water licking in the previous ST and the next nose-poke (excluding NPL>30 s). In ST, the nose-poke interval (NPI) was defined as the interval between the two rats’ nose-poking (excluding sessions with <3 ST). The water lick latency (WLL) was defined as the interval from an ST to entry into the water port. The water lick duration (WLD) was defined as the interval from entry into the water port to the last exit (excluding WLD>10 s). In 3sCo, if the NPI of an ST was <2 s or <1 s, the ST in the SR formula was replaced with the number of such trials, while the denominator remained unchanged, yielding the SR for trials meeting the <2 s or <1 s criterion.

For data shuffling, the timing of dyads was shifted by 1–100 seconds (repeated 100 times). Events in which both rats nose-poked within a given time window (3 s, 2 s, or 1 s) were counted as ST. Nose-pokes outside the specified time window were counted as FT, with multiple nose-pokes within the same time window counted as a single event. The mean SR and NPI were then calculated across the 100 shuffled datasets.

For nose approach analysis, all nose approaches occurring within 5 s before ST or FT were analyzing. For social interaction analysis (recorded by infrared detectors), all interactions occurring within 5 s before ST were extracted (SI-5s), and their total count was defined as the total interaction bout. ST that contained a social interaction within this time window were denoted as ST-SI-5s. The mean interaction duration was calculated as the total cumulative SI-5s time divided by the number of ST-SI-5s. The interaction ratio was calculated as ST-SI-5s divided by the total number of ST. The temporal distribution of SI-5s was computed in 1-s time bin.

### Video tracking and analysis based on DeepLabCut

Six body parts were selected as tracking points: the nose, left and right ears, neck, back, and tail base. To build the training set, 20 frames were extracted from each of the following sessions: day 5 of long-distance training, day 1 of 3sCo, day 1 of 2sCo, and days 1, 6, and 12 of 1sCo, across three dyads (360 frames total). Each tracking point for both rats was manually labeled in sequence using the graphical interface of maDeepLabCut (DeepLabCut Team, v.2.2.1) ^62,63^. A neural network (ResNet-50; 200 000 iterations; batch size=8) was then trained to track these body parts with high accuracy (see Figure S2A), and subsequently applied to all behavioral videos to obtain the spatial coordinates of all dyads. Missing values (NaN) were replaced by linear interpolation; tracking accuracy was defined as the proportion of frames containing valid values (non-NaN). The centroid of the triangle formed by the nose and two ears was defined as the rat’s head position.

Two regions (15×10 cm) surrounding the nose-poke port or water port were defined as the nose-poke area (NPA) or the water lick area (WLA; see Figure 2A). The interval between one rat’s head entering the NPA and its partner’s head entering the corresponding NPA was defined as the waiting time window (Figure 2B). During this period, the angle of the head-nose vector relative to the vertical line of the perforated panel was defined as α (head direction; Figure 2B). Once both rats’ heads entered the NPA, the interval before either rat initiated a nose-poking was defined as the social time window (Figure 2B). The angle of the head-nose vector relative to the partner’s head was defined as β (social orientation; Figure 2B).

Using the timing of entries into the NPA along with subsequent behavioral events (ST, FT, and nose-poking), we quantified the waiting time window and social time window for ST. In addition, we calculated the mean head direction angle (α) during the waiting time window and the mean social orientation angle (β) during the social time window. In an ST, if a waiting time window was present, that ST was defined as “waiting for partner”. In an FT, if a rat nose-poked while its partner was in the corresponding NPA, that FT was defined as “with partner in NPA”.

### Classification of cooperative strategies

Based on the durations of the waiting time window (mean value of 1sCo=1.609 s) and the social time window (mean value of 1sCo=1.648 s) observed in ST, as well as the temporal distribution of nose-poking and the interval between leaving the water lick area (ΔWLA), we categorized the cooperative strategies by which rats completed ST into three types: communication (Type 1, long social), short social (Type 2), and synchrony (Type 3), along with an unclassified category (Others). Specifically, if either rat exhibited more than 3 nose-pokes within ±1 s of the ST, or if the ST was unclassified, the trial was assigned to the “Others”. For ST with ≤3 nose-pokes, those with a social time window >1.5 s were classified as the communication strategy (Type 1). If the social time window was ≤1.5 s and both the waiting time window and the ΔWLA were ≤1.5 s, the trial was classified as the synchrony strategy (Type 2); otherwise, it was classified as the short social strategy (Type 2; social time window ≤1.5 s, and either waiting time window or ΔWLA>1.5 s).

To analyze the transition probabilities of cooperative strategies, we identified the transition sequences of the four cooperative strategies in each session (16 types of sequences in total) and calculated their frequencies of occurrence. To calculate the dominant strategy in dyads, we defined the most frequent cooperative strategy within each dyad as the dominant strategy. We then calculated the proportion of dyads for which each of these four strategies served as the dominant strategy during cooperative training. The t-distributed stochastic neighbor embedding (t-SNE) plots of sessions with different dominant strategies were calculated in MATLAB (default parameters).

### Behavioral tests for anxiety level, locomotion, and social abilities

OXT-KO rats and their WT littermates (8–9 weeks old, 300–350 g) underwent behavioral tests to assess anxiety, locomotion, social preference, and social novelty. Rats were handled for two days prior to testing. After each test, apparatuses were cleaned with 75% ethanol. All tests were video-recorded and analyzed with Noldus EthoVision XT (Noldus Information Technology, Netherlands).

The elevated plus maze (EPM) consisted of two open and two closed arms connected by a central platform. Each arm measured 50×10 cm; closed-arm walls were 40 cm high; and the maze was elevated 60 cm above the floor. Rats were placed on an open arm facing the center platform, and were recorded for 10 min. The open-field test (OFT) apparatus measured 60×60×45 cm (L×W×H), with a central zone of 30×30 cm. Rats were placed in the center, and were recorded for 15 min. The three-chamber test (TCT) apparatus measured 90×45×30 cm (L×W×H) and was divided into three equal chambers. Wire cages (18.5×13.5×20 cm, L×W×H) were placed in the side chambers, containing either a conspecific rat or a toy rat. Testing comprised three 10-min phases: habituation (both cages empty), social preference test (conspecific versus toy rat, side randomized), and social novelty test (toy replaced with a novel conspecific; not cagemate). Rats were placed in the middle chamber at the start of each phase.

### Stereotaxic surgery and microinjection

Rats were weighed and anesthetized via intraperitoneal (i.p.) injection of a mixed anesthetic (3 mL/kg for males; Zoletil 50, 5 mg/mL, Virbac, France; Xylazine, 6 mg/mL, Huamu, China). Following induction, rats were placed in a stereotaxic apparatus (Stoelting, USA) and the skin was incised to expose the skull. Craniotomy was performed at stereotaxic coordinates corresponding to the target region, with additional holes drilled for anchor screws when required. Viruses were diluted to 1–5×10^12^ viral particles/mL and loaded into glass micropipettes.

For virus-only injections (e.g., PVN expression of AAV8-hSyn-FLEX-tdTomato-T2A-Synaptophysin-EGFP; Taitool, China; Cat# S0161-8), the dura was penetrated and the viral solution was injected using a UMP3 microinjector (WPI, USA) at 80 nL/min, with ~500 nL injected per site (total dwell time ~15 min). The skin was then sutured. For optogenetic and fiber photometry experiments, anchor screws were implanted after craniotomy, followed by viral injection. Optical fibers were positioned 0.5 mm above the target site (optogenetics) or 0.2 mm above (fiber photometry), and secured with light-curing resin (Dentkist, Korea). Dental cement (mixed with carbon powder for fiber photometry) was then applied around the optical fiber and the anchor screws to reinforce the implant. Excess cement was trimmed after partial curing. After removal from the stereotaxic apparatus, all rats received gentamicin sulfate (2 mg/mL; 2.5 mL/kg, i.p.) postoperatively to prevent infection.

### Fiber photometry recording

Male SD rats (7–8 weeks old, naïve to cooperative training) received unilateral injections of AAV2/9-hSyn-OXT1.0-WPRE-hGH PA (500 nL; Brain VTA, China; Cat# PT-2741) into the piriform cortex (AP +4.2 mm, ML +2.0 mm, DV –5.8 mm) and dorsal hippocampus (AP –2.92 mm, ML +3.3 mm, DV –3.2 mm). Optical fibers (200 μm core, NA 0.5) were implanted 0.2 mm above each target. After ~1 week of recovery, behavioral training was initiated, and fiber photometry recording began ~3 weeks post-injection. OXT signals were recorded at three training stages: 3sCo, 2sCo, and the mid-phase of 1sCo (1sCo-D6). A TTL output from the AniLab controller was fed to a dual-color fiber photometry system (Thinkertech, China) for synchronization. Prior to recordings, the implanted fiber was cleaned with ethanol and connected to the patch cable; emission intensity of the 470-nm channel at the fiber’s tip was tuned to ~50 µW. About 30–50 success trials from each dyad were recorded for analysis.

For data analysis, signals were corrected for photobleaching and the isosbestic 405-nm signal was subtracted from the 470-nm channel. Then signals were aligned to behavioral events and analyzed using Z-score normalization and area under the curve (AUC). The average signals from 3s prior to ST onset (baseline) was defined as F_0_, and the relative signal change was calculated as Z-score=(F-F_0_)/STD(F_baseline_). For resampling, signals corresponding to water lick latency (WLL), water lick duration (WLD), and nose-poke latency (NPL) were resampled to 3 s (120 frames), 5 s (200 frames), and 6 s (240 frames), respectively. For each session, the OXT signal of a rat was defined as the mean across all trials (smoothed in a time window of 21 frames), and only sessions with a Z-score magnitude greater than 1.5 were included in the calculation.

### Optogenetic manipulation

Male OXT-Cre rats (7–8 weeks old, naïve to cooperative training) received bilateral PVN (AP –1.8 mm, ML +1.5 mm, DV –7.9 mm, 7.5° angle) injections of AAV2/9-hSyn-DIO-stGtACR2-eGFP-WPRE-pA (500 nL; Taitool, China; Cat# S0625-9) or control AAV2/9-hSyn-DIO-EGFP-WPRE-pA (500 nL; Taitool, China; Cat# S0746-9). Optical fibers (200 μm core, NA 0.37) were implanted 0.5 mm above the PVN. After ~1 week of recovery, behavioral training was initiated, and optogenetic manipulation began ~3 weeks post-injection (from 3sCo to 1sCo; 18 days total). SLOC Laser (Shanghai Laser & Optics Century, China) was controlled via TTL from the AniLab controller. During cooperative training, a blue light (473 nm, 8–12 mW, ≤15 s) was delivered from ST onset to water-lick end (WLL-WLD).

For simultaneous PVN manipulation and OXT recording in the dHPC or Pir, bilateral PVN expression of AAV2/9-hSyn-DIO-hChR2(H134R)-mCherry-ER2-WPRE-pA (500 nL; Taitool, China; Cat# S0312-9) or AAV2/9-hSyn-DIO-stGtACR2-eGFP-WPRE-pA (500 nL; Taitool, China; Cat# S0625-9) was combined with unilateral dHPC/Pir expression of AAV2/9-hSyn-OXT1.0-WPRE-hGH PA (500 nL; Brain VTA, China; Cat# PT-2741), and matched optical fibers were implanted. Optogenetic activation (PVN, bilateral) used a 12 mW blue light at 20 Hz, 30% duty cycle, 10 s, ITI 1 min; optogenetic inhibition (PVN, bilateral) used an 8–12 mW blue light, 10 s, ITI 1 min. A TTL from the AniLab controller (also control SLOC Laser) was fed to the fiber photometry system for synchronization.

### Whole-cell patch clamp electrophysiology

To validate the efficacy of stGtACR2 in PVN OXT-ergic neurons, *ex vivo* whole-cell patch-clamp recordings were performed in brain slices from OXT-Cre rats expressing AAV2/9-hSyn-DIO-stGtACR2-eGFP (500 nL; Taitool, China; Cat# S0625-9) for ~3 weeks. The detailed protocol has been described previously ^104,105^. Briefly, brain slices (350 μm) were prepared in ice-cold oxygenated slicing solution, incubated at 32℃ for 12 min, and then maintained in room-temperature artificial cerebrospinal fluid (ACSF). Recordings were obtained with 3–7 MΩ glass microelectrodes under current clamp, with series resistance compensated and only cells with Ra<25 MΩ and Ih<100 pA were analyzed. eGFP+ cells were held near –50 mV to elicit stable firing, and blue light pulses (470 nm, 500 or 1 000 ms) were applied to assess the effect on membrane potential and action potential firing.

### Immunohistochemistry and imaging

Rats were sacrificed under anesthesia and perfused transcardially with phosphate-buffered saline (PBS), followed by 4% paraformaldehyde (PFA). Brains were then removed and dehydrated in 30% sucrose at 4℃. Coronal sections (30–50 μm) containing the target regions (e.g., PVN, Pir, dHPC) were cut on a CM1950 cryostat (Leica, Germany) and mounted on glass slides. For immunohistochemistry, rabbit anti-oxytocin (1:10 000; Phoenix Biotech, USA; Cat# G-051-01) was used as the primary antibody, and donkey anti-rabbit Alexa Fluor 555 (1:500; Thermo Fisher, USA; Cat# A-21206) was used as the secondary antibody. After secondary antibody incubation, brain slices were mounted on glass slides. All slices were counterstained with DAPI Fluoromount-G (Southern Biotech, China; Cat# 0100-20) and then imaged under an Olympus VS120 fluorescence microscope with a 10× air objective.

### Statistics

The data were analyzed using Prism 10 (GraphPad Software, USA) and MATLAB (MathWorks, USA). Specific statistical analyses are indicated in the figure legends for each figure and primarily include paired or unpaired t-tests, Mann–Whitney tests, Wilcoxon signed-rank tests, F-tests, and one-way or two-way analyses of variance (ANOVA). Normality was assessed using the Shapiro–Wilk or Kolmogorov–Smirnov tests, followed by F-tests for homogeneity of variance. Symbols *, **, and *** indicate *P*< 0.05, *P*< 0.01, and *P*<0.001, respectively, while “ns” denotes no significant difference.

## AUTHORS’ CONTRIBUTIONS

Y. L. and Z. W. designed the experiments and wrote the paper. Y. L. performed the behavioral experiments and data analysis. L. W. performed the electrophysiological recording and analysis. Q. W. and Y. L. bred and genotyped OXT-KO and OXT-Cre rats. All authors read and approved the final version of the manuscript.

## ACKNOWLEDGEMENTS

We thank Shangyi Wang for assistance with writing Matlab code and Xintong Zhou for assistance with installing DeepLabCut. We are also grateful to technical support from the optical imaging facility and animal facility of the Institute of Neuroscience, Center for Brain Science and Intelligence Technology, CAS.

## Supplementary Materials

**Supplementary Figure S1.**
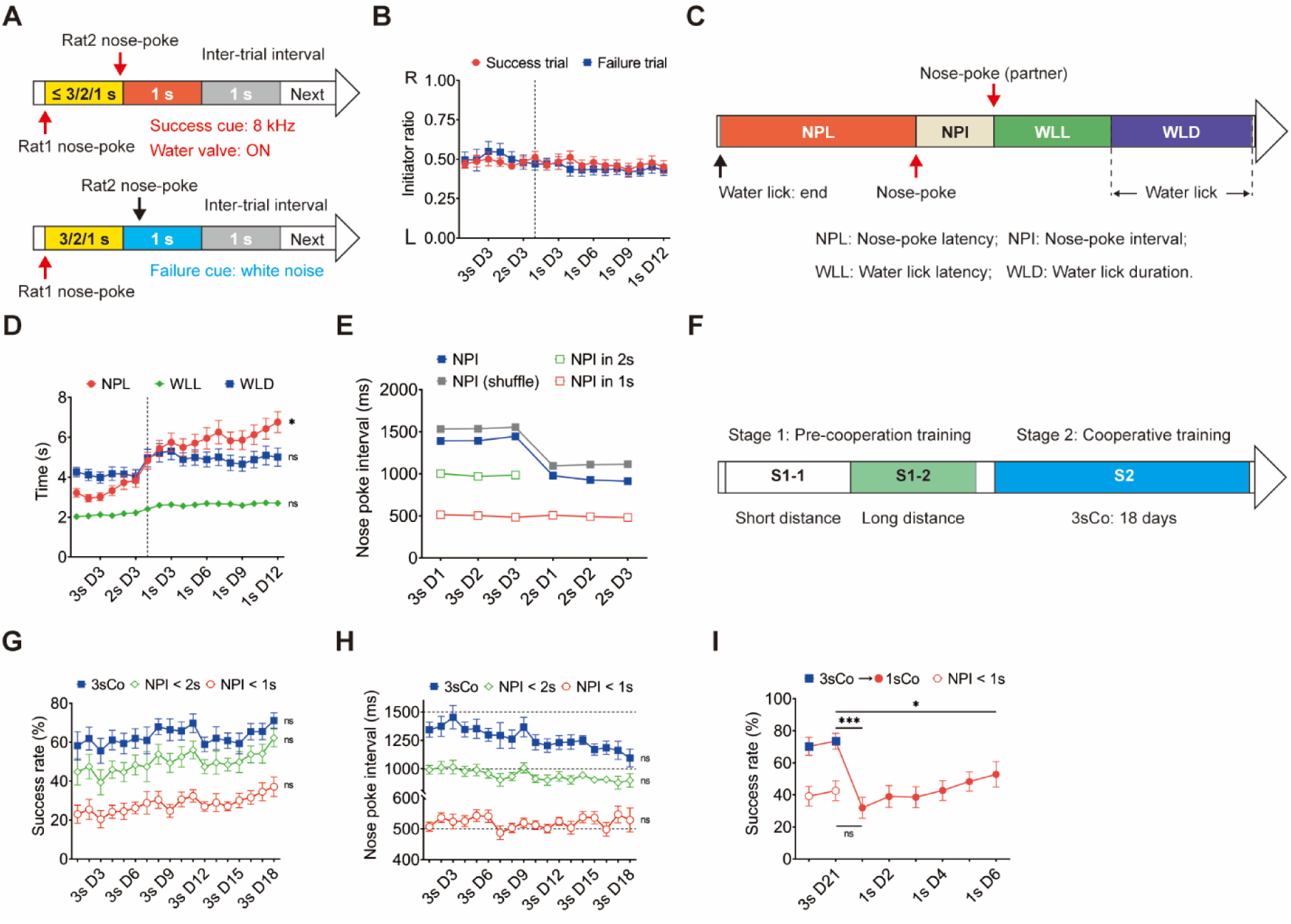
Behavioral performance of dyads during 1sCo and 3sCo training. A: Diagram of the success trial (ST, top) and the failure trial (FT, bottom). Yellow box is the cooperation time window (3s, 2s, or 1s); red or blue box is the period that success or failure cues are delivered; grey box is the inter-trial interval. B: Initiator ratios of success trial and failure trial for dyads during cooperative training (*n*=20). C: Diagram of four behavioral events: nose-poke latency (NPL), nose-poke interval (NPI), water lick latency (WLL), and water lick duration (WLD). D: NPL (red), WLL (green), and WLD (blue) of dyads during cooperative training (*n*=20; 1sCo: NPL, *P*=0.0143; WLL, *P*=0.1414; WLD, *P*=0.3006). E: NPI of dyads during 3sCo and 2sCo training (blue line, *n*=20). The gray line indicates the shuffled control of NPI; green and red lines indicate the NPI within 2 s and 1 s, respectively. F: Diagram of the training process of 18-day 3sCo task. G: Success rate (SR) of dyads during 3sCo training (blue line, *n*=8, *P*=0.3829). Green and red lines indicate the SR with NPI values within 2 s and 1 s, respectively (NPI<2s, *P*=0.2791; NPI<1s, *P*=0.2572). H: NPI of dyads during 3sCo training (blue line, *n*=8, *P*=0.1635). Green and red lines indicate the NPI within 2 s and 1 s, respectively (NPI<2s, *P*=0.3191; NPI<1s, *P*=0.4891). I: SR of dyads that directly transferred from 3sCo to 1sCo (*n*=8; 3s D21 vs. 1s D1, *P*=0.0001; 3s D21 vs. 1s D6, *P*=0.0134). Open dots indicate the SR which NPI values within 1 s (3s D21 vs. 1s D1, *P*=0.0735). In B and D, vertical dashed lines indicate D1 of 1sCo. Data are presented as mean ± SEM. Statistics: repeated measures one-way ANOVA (D and G–I). ns: Not significant; *: *P*<0.05; **: *P*<0.01; ***: *P*<0.001.

**Supplementary Figure S2.**
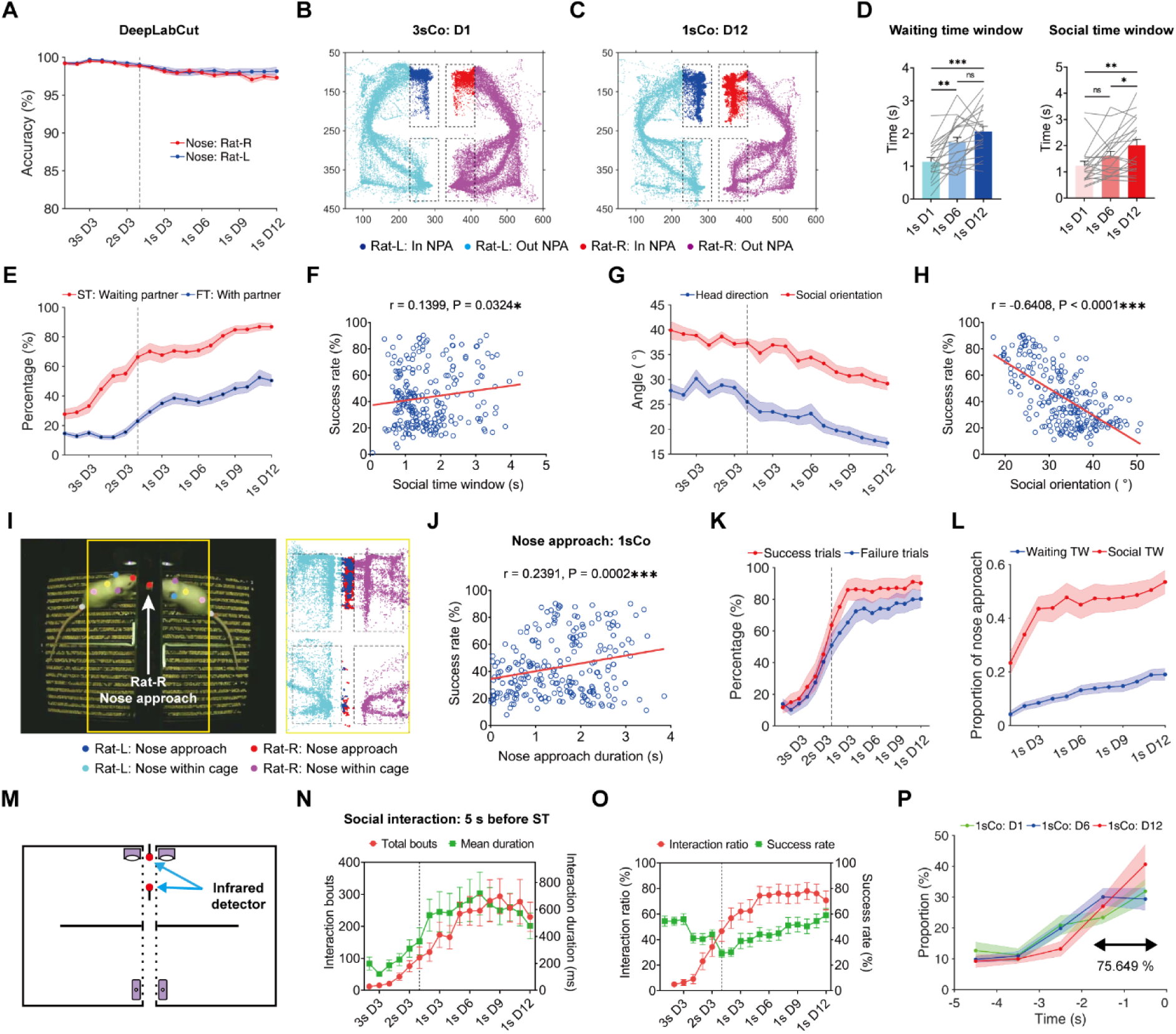
Analysis of social interaction during cooperative training. A: Tracking accuracy of dyadic noses using DeepLabCut (*n*=20). B–C: Example trajectories of dyadic heads during 3sCo-D1 (B) and 1sCo-D12 (C). Axes represent video pixel coordinates; dark coloring dots indicate head positions within the nose-poke area (NPA, upper dashed boxes). D: Comparisons of waiting time window (left) and social time window (right) for dyads among the D1, D6, and D12 of 1sCo (*n*=20). Waiting time window: D1 vs. D6, *P*=0.0011; D1 vs. D12, *P*=0.0001; D6 vs. D12, *P*=0.1081. Social time window: D1 vs. D6, *P*=0.0756; D1 vs. D12, *P*=0.0050; D6 vs. D12, *P*=0.0230. E: Changes of the percentages that dyads wait for partner in success trials (ST, waiting for partner) and failure trials (FT, with partner in NPA) during cooperative training (*n*=20). F: Linear regression of the duration of social time window versus success rate (SR) during 1sCo (all sessions, *n*=234, *P*=0.0324). G: Changes in head direction and social orientation of dyads during cooperative training (*n*=20). H: Linear regression of social orientation versus SR during 1sCo (all sessions, *n*=234, *P*<0.0001). I: Schematic of the nose approach during cooperation (left) and example trajectories of dyadic noses during 1sCo-D12 (right). Dark coloring dots indicate nose positions across rectangular perforations. J: Linear regression of nose approach duration versus SR during 1sCo (all sessions, *n*=234, *P*=0.0002). K: Percentage of trials with nose approach (>100 ms) for success trials (red) and failure trials (blue) during cooperative training (*n*=20). L: Proportion of time spent in nose approach during waiting time window (blue) and social time window (red, *n*=20). M: Diagram of the infrared detector positioned in the middle of cages. N: Total bouts (red) and mean duration (green) of social interaction for dyads during cooperative training (*n*=11). O: Social interaction ratio (red) and success rate (green) of dyads during cooperative training (*n*=11). P: Distribution of social interactions within the 5 seconds before successful cooperation during 1sCo (*n*=11); 75.649% for D12 of 1sCo. In A, E, G, K and N–O, vertical dashed lines indicate D1 of 1sCo. Data are presented as mean ± SEM. Statistics: repeated measures one-way ANOVA (D) and F-test (F, H and J). ns: Not significant; *: *P*<0.05; **: *P*<0.01; ***: *P*<0.001.

**Supplementary Figure S3.**
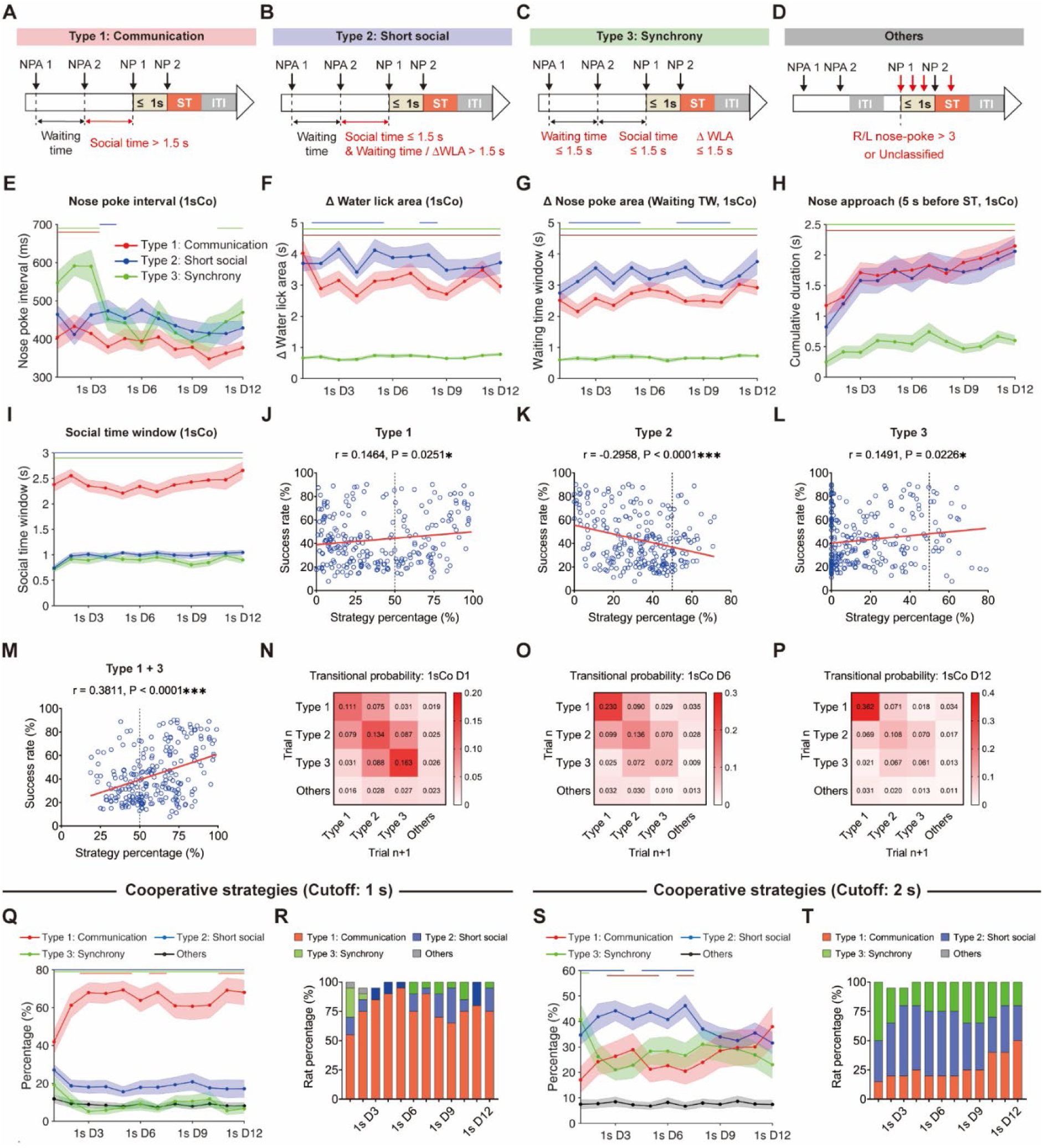
Cooperative strategies of dyads during cooperation. A–D: Schematics of different types of cooperative strategies: communication (type 1, A), short social (type 2, B), synchrony (type 3, C), and others (D). NP, NPA, ST, and ITI denote nose-poke, nose-poke area, success trial, and inter-trial interval, respectively. ΔWLA indicates the interval between two rats leaving the WLA. E: Changes in the nose poke interval of three cooperative strategies during 1sCo phase (*n*=20). F–G: Changes in the interval between leaving the water lick area (ΔWLA, F) and the interval between leaving the nose poke area (ΔNPA or waiting time window, G) of three cooperative strategies during 1sCo phase (*n*=20). H–I: Changes in the duration of nose approach (H) and the social time window (I) of three cooperative strategies during 1sCo phase (*n*=20). J–M: Linear regressions of strategy percentage versus success rate during 1sCo (all sessions, *n*=234; J, Type 1, *P*=0.0251; K, Type 2, *P*<0.0001; L, Type 3, *P*=0.0226; M, Type 1+3, *P*<0.0001). Vertical dashed lines indicate 50%. N–P: Transitional probabilities of strategy types from the present to the subsequent success trial in the D1 (N), D6 (O), and D12 (P) of 1sCo (*n*=20). Q: Percentage changes in different cooperative strategies (cutoff: 1 s) during 1sCo phase (*n*=20). R: Percentage of dyads using the dominant strategies (cutoff: 1 s) during 1sCo phase. S: Percentage changes in different cooperative strategies (cutoff: 2 s) during 1sCo phase (*n*=20). T: Percentage of dyads using the dominant strategies (cutoff: 2 s) during 1sCo phase. In E–I, Q and S, lines at the top indicate statistical significance (*P*<0.05): Type 1 vs. Type 2 (blue), Type 1 vs. Type 3 (green), and Type 2 vs. Type 3 (red). Data are presented as mean ± SEM. Statistics: one-way ANOVA (E–I, Q and S) and F-test (J–M). *: *P*<0.05; **: *P*<0.01; ***: *P*<0.001.

**Supplementary Figure S4.**
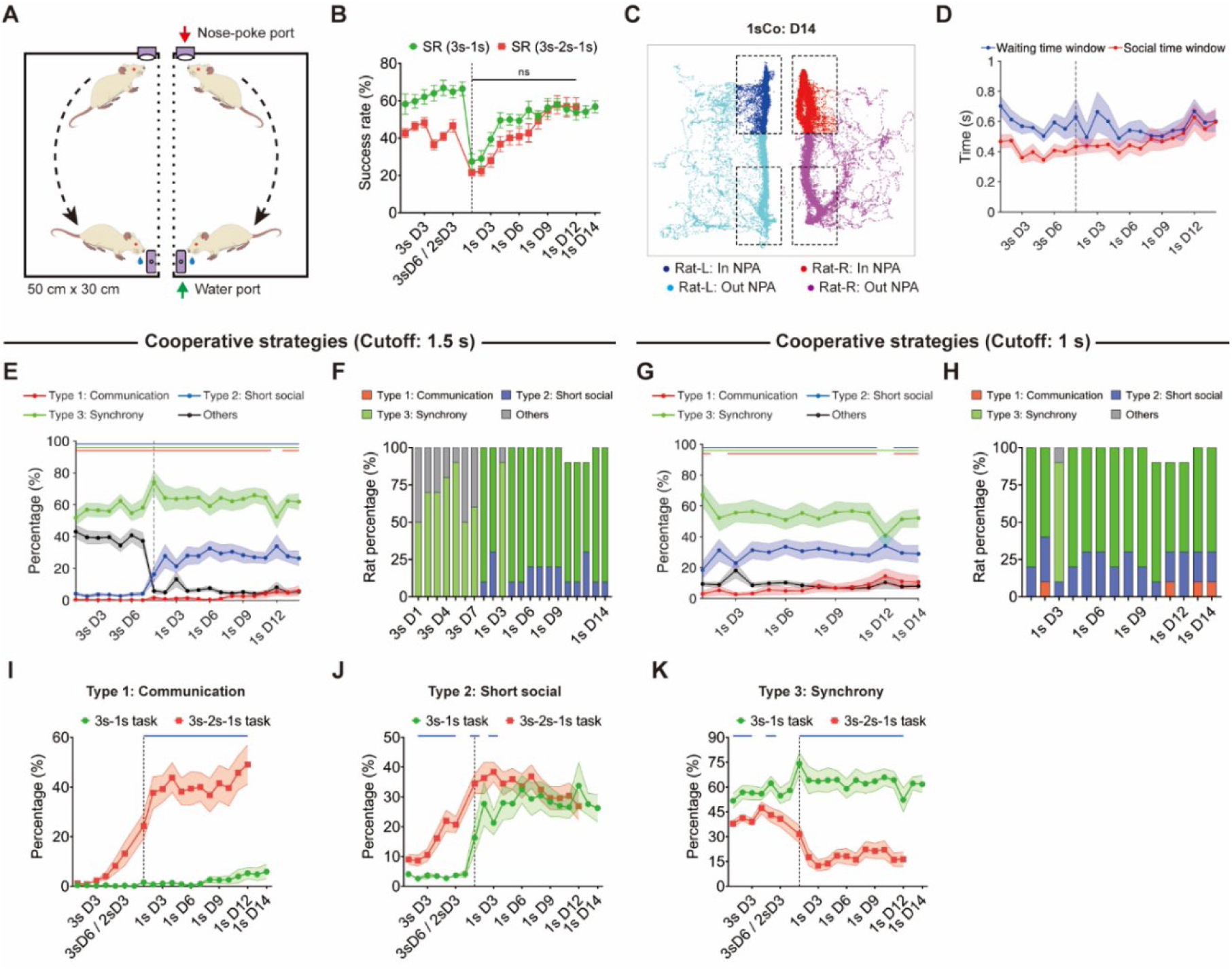
Classification of cooperative strategies during 3sCo-1sCo training. A: Schematic of the behavioral apparatus and task process. Each cage contains a nose-poke port (red arrow) and a water port (green arrow), with middle dashed lines indicating the rectangular perforations. B: Success rate (SR) of 3sCo-1sCo dyads (green, *n*=10) and 3sCo-2sCo-1sCo dyads (red, *n*=20) during cooperative training. C: Example trajectories of dyadic heads during 1sCo-D14. Dark coloring dots indicate head positions within the nose-poke area (NPA, upper dashed boxes). D: Changes in the waiting time window (blue) and social time window (red) of dyads during cooperative training (*n*=10). E: Percentage changes in different cooperative strategies (cutoff: 1.5 s) during cooperative training (*n*=10). F: Percentage of dyads using the dominant strategies (cutoff: 1.5 s) during cooperative training. G: Percentage changes in different cooperative strategies (cutoff: 1 s) during 1sCo phase (*n*=10). H: Percentage of dyads using the dominant strategies (cutoff: 1 s) during 1sCo phase. I–K: Percentage of strategy Type 1 (I), Type 2 (J) and Type 3 (K) in 3sCo-2sCo-1sCo task (green, *n*=10) and 3sCo-2sCo-1sCo task (red, *n*=20). In B, D–E and I–K, vertical dashed lines indicate D1 of 1sCo. In E and G, lines at the top indicate statistical significance (*P*<0.05): Type 1 vs. Type 2 (blue), Type 1 vs. Type 3 (green), and Type 2 vs. Type 3 (red). In I–K, the blue lines at the top indicate statistical significance (*P*<0.05). Data are presented as mean ± SEM. Statistics: two-tailed unpaired *t*-test (B) and one-way ANOVA (E, G and I–K).

**Supplementary Figure S5.**
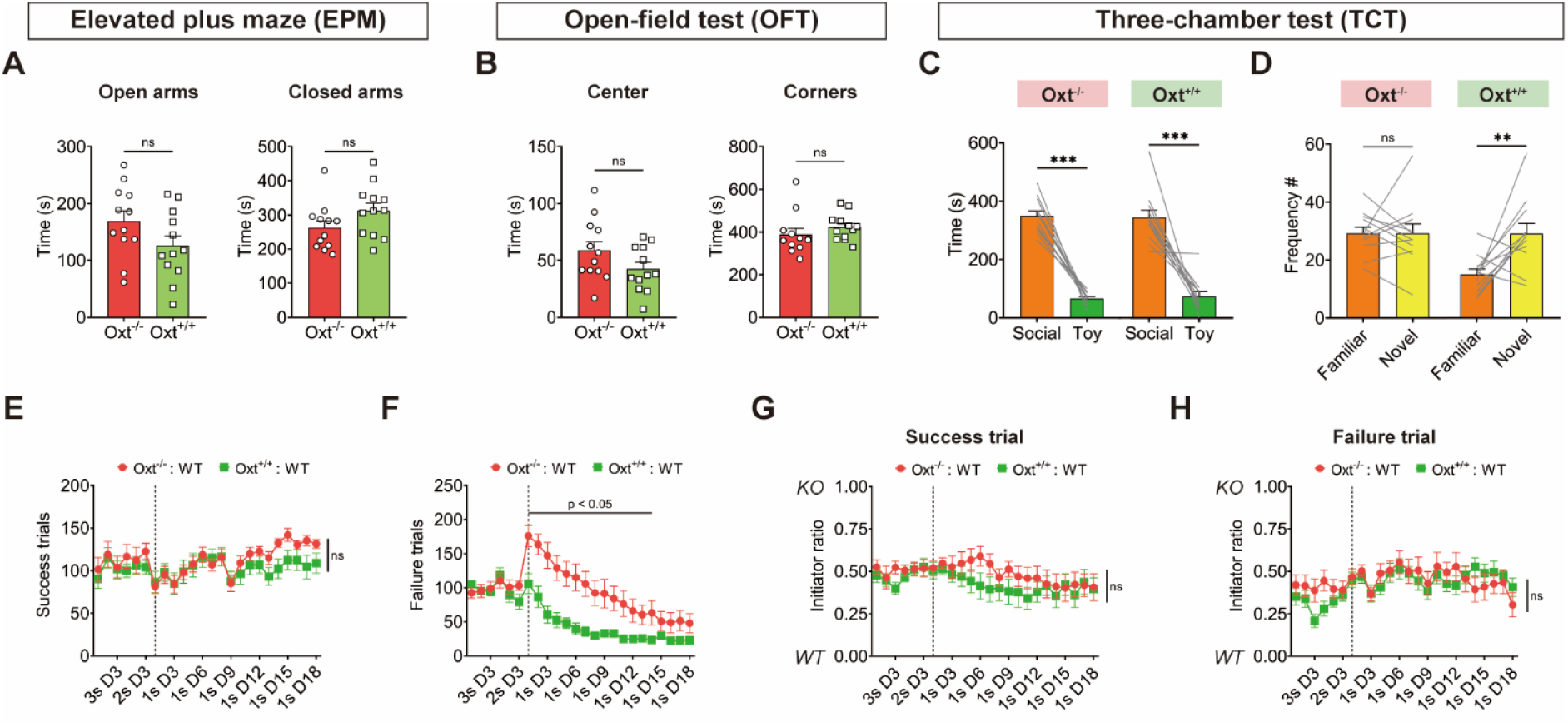
Behavioral phenotypes of OXT knockout rats during tests and cooperative training. A: Durations of OXT-KO rats and their WT littermates (WT-LM) in open arms (left, *P*=0.0978) and closed arms (right, *P*=0.1028) during the elevated plus maze test (EPM, *n*=12 and 12). B: Durations of OXT-KO and WT-LM rats in center area (left, *P*=0.1065) and corners (right, *P*=0.0684) during the open-field test (OFT, *n*=12 and 12). C: Time spent on social versus toy sides of OXT-KO and WT-LM rats during the three-chamber test (TCT, *n*=12 and 12; OXT-KO, *P*<0.001; WT-LM, *P*<0.001). D: Frequency of entries to familiar versus novel sides of OXT-KO and WT-LM rats during TCT (*n*=12 and 12; OXT-KO, *P*>0.9999; WT-LM, *P*=0.0088). E–F: Success trials (E, 1s D1–D18, *P*=0.4533) and failure trials (F) of OXT-KO and WT-LM dyads during cooperative training (*n*=11 and 13). G–H: Initiator ratios of OXT-KO and WT-LM dyads in success trials (G, 1s D1–D18, *P*=0.4039) and failure trials (H, 1s D1–D18, *P*=0.9354) during cooperative training (*n*=11 and 13). In E–H, vertical dashed lines indicate D1 of 1sCo. Data presented as mean ± SEM. Statistics: unpaired *t*-tests (A, B-left, and F), paired *t*-tests (C–D), Mann Whitney test (B-right), and two-way ANOVA (E and G–H). All statistical tests are two-tailed (except for two-way ANOVA). ns: Not significant; *: *P*<0.05; **: *P*<0.01; ***: *P*<0.001.

**Supplementary Figure S6.**
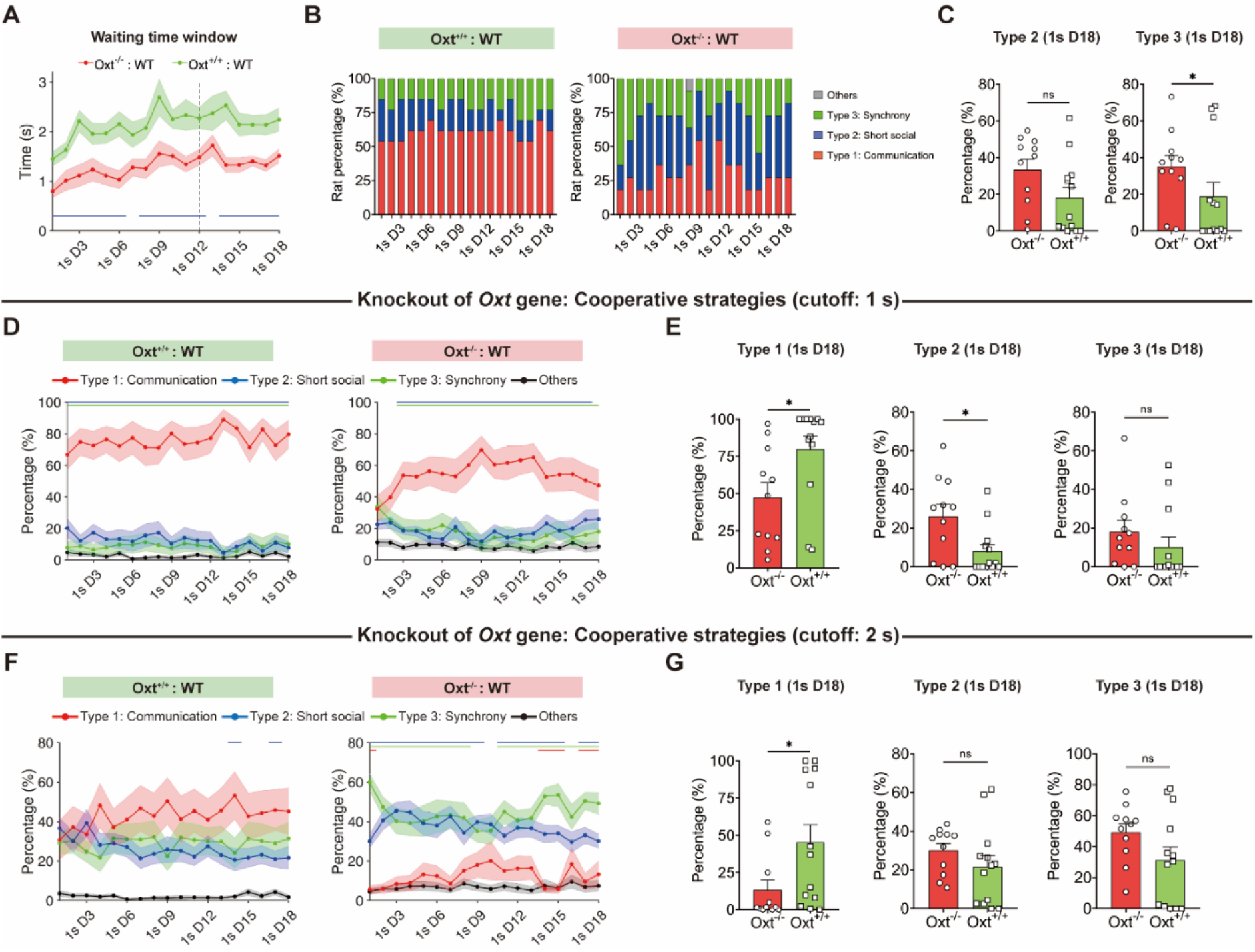
Formation of cooperative strategies in OXT knockout rats. A: Durations of the waiting time window in OXT-KO and WT littermate (WT-LM) dyads during 1sCo training (*n*=11 and 13). Vertical dashed line indicates D12 of 1sCo. B: Percentage of WT-LM (left) and OXT-KO dyads (right) that used the dominant cooperative strategies during 1sCo training. C: Percentages of strategy Type 2 (short social) and Type 3 (synchrony) in OXT-KO and WT-LM dyads on D18 of 1sCo (*n*=11 and 13; Type 2, *P*=0.0718; Type 3, *P*=0.0465). D: Percentage changes of three cooperative strategies (cutoff: 1 s) in WT-LM (left, *n*=13) and OXT-KO dyads (right, *n*=11) during 1sCo training. E: Percentage of strategy Type 1, Type 2 and Type 3 in OXT-KO and WT-LM dyads on D18 of 1sCo (*n*=11 and 13; Type 1: *P*=0.0126, Type 2: *P*=0.0263, Type 3: *P*=0.0781). F: Percentage changes of three cooperative strategies (cutoff: 2 s) in WT-LM (left, *n*=13) and OXT-KO dyads (right, *n*=11) during 1sCo training. G: Percentage of strategy Type 1, Type 2 and Type 3 in OXT-KO and WT-LM dyads on D18 of 1sCo (*n*=11 and 13; Type 1: *P*=0.0311, Type 2: *P*=0.2445, Type 3: *P*=0.1053). In A, lines at bottom indicate statistical significance (*P*<0.05). In D and F, lines at the top indicate statistical significance (*P*<0.05): Type 1 vs. Type 2 (blue), Type 1 vs. Type 3 (green), and Type 2 vs. Type 3 (red). Data are presented as mean ± SEM. Statistics: one-way ANOVA (A, D and F) and two-tailed Mann Whitney test (C, E and G). ns: Not significant; *: *P*<0.05.

**Supplementary Figure S7.**
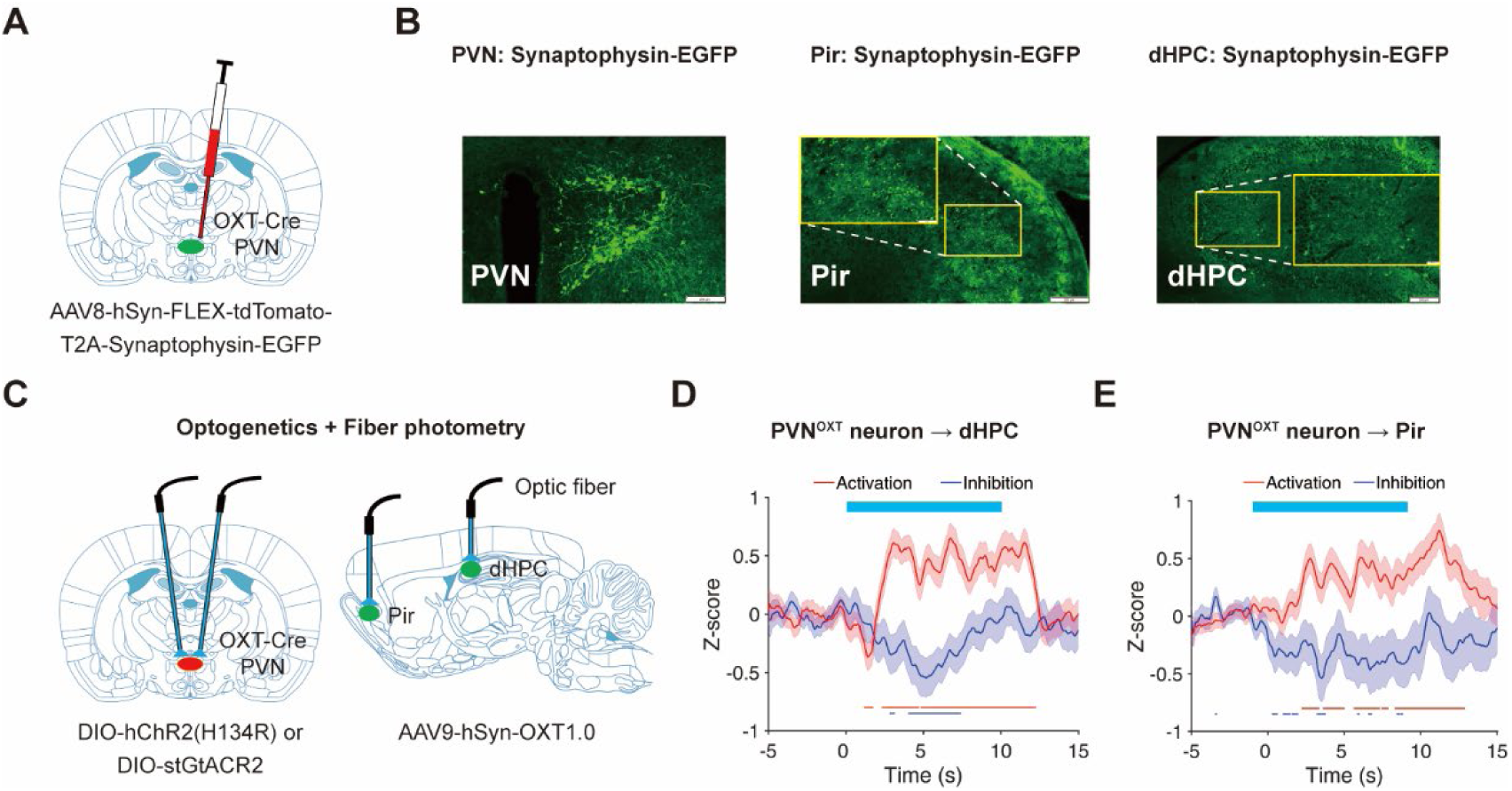
PVN OXT-ergic neurons mediate OXT release in dHPC and Pir. A: Synaptophysin-EGFP expression in the PVN of OXT-Cre rats. B: Representative Synaptophysin-EGFP expression in the PVN, Pir, and dHPC of OXT-Cre rats. Scale bars: 200 μm (large image), 50 μm (inset). C: DIO-hChR2 or DIO-stGtACR2 expressed in the PVN for optogenetic manipulation; OXT1.0 expressed in the Pir and dHPC for fiber photometry recording (OXT-Cre rats). D–E: OXT signal changes in the dHPC (D, activation: 2 rats, 71 trials; inhibition: 2 rats, 76 trials) and Pir (E, activation: 3 rats, 115 trials; inhibition: 2 rats, 77 trials) during optogenetic activation/inhibition of PVN OXT-ergic neurons (10 s). Lines at bottom: activation versus baseline (red) and inhibition versus baseline (blue). In D–E, lines at bottom indicate statistical significance (*P*<0.05). Data are presented as mean ± SEM. Statistics: one-way ANOVA (D–E).

**Supplementary Figure S8.**
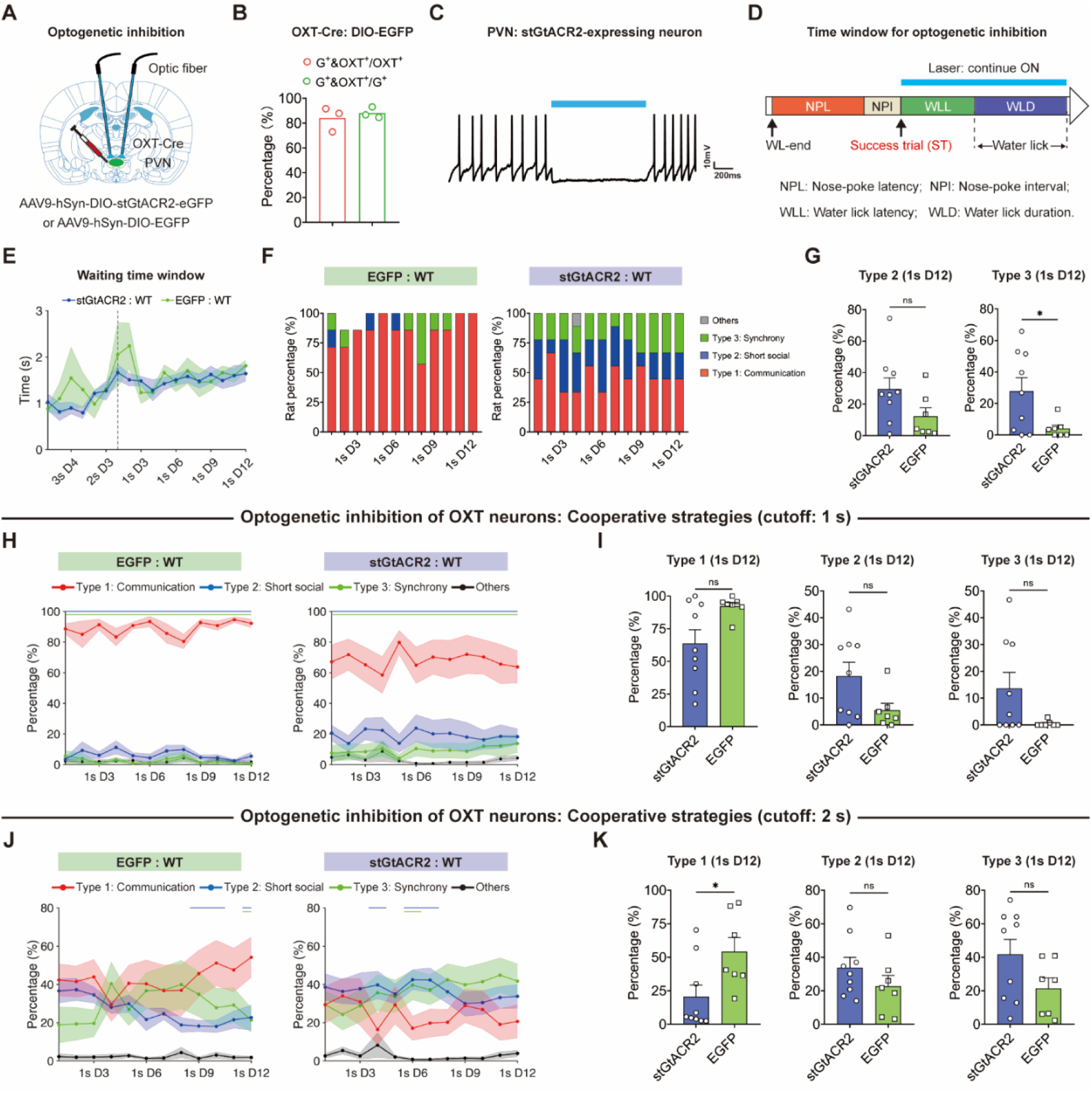
Formation of cooperative strategies when inhibiting PVN OXT-ergic neurons during learning. A: Bilateral expression of stGtACR2 or EGFP in the PVN of OXT-Cre rats for optogenetic inhibition. B: Expression efficiency of DIO-EGFP in the PVN of OXT-Cre rats. G+ represents EGFP-positive, and OXT+ represents OXT-positive. C: Blue light stimulation (1000 ms) could inhibit the excitability of stGtACR2-expressing neuron in the PVN in *ex vivo* brain slices. D: Schematic of the time window (WLL-WLD) for optogenetic inhibition. Abbreviation: nose-poke latency (NPL), water lick latency (WLL), water lick duration (WLD). E: Durations of the waiting time window in stGtACR2 and control dyads during cooperative training (*n*=9 and 7). Vertical dashed line indicates D1 of 1sCo. F: Percentage of control dyads (left) and stGtACR2 dyads (right) that used the dominant cooperative strategies during 1sCo training. Gaps indicate missing behavioral videos. G: Percentages of strategy Type 2 (short social) and Type 3 (synchrony) in stGtACR2 and control dyads on D12 of 1sCo (*n*=9 and 7; Type 2, *P*=0.0870; Type 3, *P*=0.0316). H: Percentage changes of three cooperative strategies (cutoff: 1 s) in control dyads (left, *n*=7) and stGtACR2 dyads (right, *n*=9) during 1sCo training. I: Percentage of strategy Type 1, Type 2 and Type 3 in stGtACR2 and control dyads on D12 of 1sCo (*n*=9 and 7; Type 1: *P*=0.1188, Type 2: *P*=0.1188, Type 3: *P*=0.1320). J: Percentage changes of three cooperative strategies (cutoff: 2 s) in control dyads (left, *n*=7) and stGtACR2 dyads (right, *n*=9) during 1sCo training. K: Percentage of strategy Type 1, Type 2 and Type 3 in stGtACR2 and control dyads on D12 of 1sCo (*n*=9 and 7; Type 1: *P*=0.0229, Type 2: *P*=0.2374, Type 3: *P*=0.1026). In H and J, lines at the top indicate statistical significance (*P*<0.05): Type 1 vs. Type 2 (blue), Type 1 vs. Type 3 (green), and Type 2 vs. Type 3 (red). Data are presented as mean ± SEM. Statistics: one-way ANOVA (E, H and J) and two-tailed unpaired *t*-test (G, I and K). ns: Not significant; *: *P*<0.05.

